# Genetic silencing of AKT induces melanoma cell death

**DOI:** 10.1101/2022.08.15.504039

**Authors:** Gennie L. Parkman, Tursun Turapov, David A. Kircher, William J. Burnett, Christopher M. Stehn, Kayla O’Toole, Ryan Flaherty, Riley C. Elmer, Katie M. Culver, Mona Foth, Robert H. I. Andtbacka, David H. Lum, Robert Judson-Torres, Matthew W. VanBrocklin, Sheri L. Holmen, Martin McMahon

**Affiliations:** Huntsman Cancer Institute, University of Utah Health Sciences Center, Salt Lake City, Utah 84112, USA; Department of Oncological Sciences, University of Utah Health Sciences Center, Salt Lake City, Utah 84112, USA; Department of Dermatology, University of Utah Health Sciences Center, Salt Lake City, Utah 84112, USA; Department of Surgery, University of Utah Health Sciences Center, Salt Lake City, Utah 84112, USA

**Author notes:** **Corresponding Author:** Sheri L. Holmen, PhD. Huntsman Cancer Institute, University of Utah, 2000 Circle of Hope Salt Lake City, Utah 84112. **Declaration of interests:** The authors declare no potential conflicts of interest.

**Keywords:** AKT, SGK, melanoma

## Abstract

Aberrant activation of the PI3K-AKT pathway is common in melanoma but efforts to drug this pathway have proven largely ineffective in patients. In this study, we observed that pharmacological inhibition of AKT was ineffective whereas genetic silencing of all three AKT paralogs significantly abrogated melanoma cell growth through effects on mTORC signaling. This phenotype could be rescued by overexpression of AKT but was dependent on kinase activity. Interestingly, expression of the serine/threonine kinase SGK1 was increased following genetic suppression of AKT but only expression of activated SGK1 could rescue the lethal effect of AKT knockdown. SGK1 also increases tumor growth and decreases survival in a BRAF^V600E^-driven mouse model of melanoma. Pharmacological inhibition of SGK and AKT reduced cell proliferation. These results demonstrate that SGK1 can compensate for AKT loss in this context and suggest that dual targeting of SGK1 and AKT may represent a novel therapeutic strategy in this disease.

**SIGNIFICANCE:** Although the AKT1, 2 & 3 genes encode: *Bona fide* oncoprotein kinases; well-validated downstream effectors of PI3’-lipids and: pleiotropic regulators of numerous aberrant properties of cancer cells, their role in melanoma progression and/or maintenance is poorly understood. Here we explore the effects of genetic or pharmacological inhibitors of AKT1-3 and conclude that combined inhibition of AKT plus the related SGK protein kinases may be required to inhibit melanomagenesis.

## INTRODUCTION

Approximately fifty-percent of all melanomas harbor a *BRAF^T1799A^* mutation that encodes for the BRAF^V600E^ oncoprotein kinase, the expression of which leads to constitutive activation of MAP kinase signaling (1). The importance of this pathway in melanoma maintenance is emphasized by FDA approval of drugs that target BRAF^V600E^ plus MEK for the treatment of *BRAF*-mutated melanoma (2). However, in both preclinical models and in human melanocytes, expression of BRAF^V600E^ drives the development of benign melanocytic neoplasia that generally fail to progress to melanoma without additional cooperating alterations (3). While the mechanism(s) by which BRAF^V600E^ cooperates with other alterations to convert normal melanocytes to metastatic melanoma cells has yet to be fully elucidated (4,5), aberrant activation of phosphatidylinositol 3’-lipid (PI3’-lipid) signaling drives progression of BRAF^V600E^-driven melanoma in genetically engineered mouse (GEM) models (6). Moreover, constitutive activation of PI3’-lipid signaling is observed widely across human malignancies, including melanoma, either due to silencing of the PTEN PI3’-lipid phosphatase tumor suppressor or mutational activation of *PIK3C* genes encoding PI3’-kinases (e.g. *PIK3CA* encoding PI3’-kinase-α). Of the various known effectors of PI3’-lipid signaling, the AKT (AKT1, 2 & 3) family of protein kinases are thought to play a prominent role in melanomagenesis (7). In normal cells, AKT protein kinases are activated downstream of numerous growth factor receptors that activate PI3’-lipid signaling. However, in cancer cells, activating mutations or copy number gains of *AKT* genes have been reported in melanoma patient samples (8). The three highly homologous AKT paralogs normally reside in the cytoplasm in an inactive conformation. However, in response to increased activity of PI3’-kinases, they bind via their N-terminal pleckstrin homology (PH) domains to PI3’-lipids at the cell membrane where they undergo activating phosphorylation by PDK1 and mTORC2 (9,10). Once activated, AKT protein kinases are reported to phosphorylate a multitude of target proteins that, in turn, regulate cellular survival, growth, motility, and invasion (11,12). In melanoma, the level of phospho-AKT is reported to steadily increase throughout progression from dysplastic nevi to metastatic melanomas. Moreover, silencing of PTEN or mutational activation of AKT are reported as mechanisms of acquired resistance to FDA-approved inhibitors of BRAF^V600E^ signaling in *BRAF*-mutated melanoma, highlighting the potential importance of targeting PI3K>AKT signaling for the treatment of this disease (13).

In preclinical models, inhibition of PI3K signaling has a cytostatic effect on melanoma cell proliferation. In contrast, pharmacological inhibition of AKT, including the use of structurally unrelated and mechanistically dissimilar agents, has little to no effect on melanoma cell proliferation (14). Furthermore, clinical trials evaluating AKT inhibitors as single agents have also had minimal efficacy in melanoma, suggesting that AKT may not be essential for melanoma maintenance (15). However, this conclusion is complicated by negative feedback loops mediated by mTORC1 and/or p70^S6K^ that can lead to activation of the PI3K signaling pathway following AKT inhibition (16). Our previous research indicated that the effects of PI3K inhibitors on the activity of mTORC1 leading to regulation of protein synthesis is independent of AKT (17). Moreover, additional PI3K-dependent protein kinases, such as the serum and glucocorticoid-regulated kinases (SGK) 1-3 family, have been identified and are known to have similar downstream effectors as AKT (18). Finally, although PI3’-kinase inhibition forestalled the onset of MEK inhibitor resistance in BRAF^V600E^-driven melanoma (14), translation of such observations to routine clinical use has been hampered by the attendant toxicity of combined inhibition of BRAF^V600E^ plus PI3’-kinase signaling in patients (19,20). Hence, these data highlight the critical need to understand the importance of PI3K>AKT signaling in melanoma maintenance as a step towards possible combination pathway-targeted therapy in the subset of melanomas in which these pathways cooperate for melanoma proliferation and survival.

Here, we evaluated the role of AKT in melanoma cell proliferation and observed a striking difference in melanoma cell viability between pharmacological inhibition versus RNAi-mediated (siAKT1-3) silencing of total AKT expression. We noted that RNAi-mediated inhibition of AKT1-3 expression resulted in striking melanoma cell lethality that was rescued by expression of siAKT1-3 resistant forms of AKT. Interestingly, we observed that a combination of three siRNAs against AKT1, 2, and 3 (henceforth referred to as siAKT1-3) significantly decreased mTORC activity whereas pharmacological inhibition of AKT did not. Furthermore, catalytically active SGK was also able to rescue the effects of siAKT1-3. Pharmacological inhibition of both AKT and SGK led to a strikingly superior decrease in melanoma cell proliferation compared with targeting either protein alone and this was mediated by suppression of mTORC signaling. These data serve to emphasize the complexity of PI3’-lipid signaling in melanoma progression and maintenance and firmly indicate that conventional pharmacological blockade of AKT is unlikely to be a successful strategy for the treatment of *BRAF*-mutated melanoma.

## RESULTS

### siRNA-mediated knockdown of all three AKT paralogs (siAKT1-3) is lethal to melanoma cells

There is an abundance of evidence suggesting that AKT plays a crucial and central role in tumorigenesis, particularly melanomagenesis such that *AKT1, 2 & 3:* Encode *bona fide* oncoprotein kinases; are well-validated downstream effectors of PI3’-lipids and: AKT substrates are pleiotropic regulators of numerous aberrant properties of the cancer cell (21). However, their role in melanoma progression and maintenance is poorly understood and, since AKT inhibitors have largely failed in melanoma clinical trials (22). Therefore, we first sought to test the possible importance of AKT for melanoma cell proliferation *in vitro* using multiple complementary approaches. Our previous data indicated that pharmacological AKT inhibitors had little or no effect on melanoma cell proliferation despite confirmed effects on phosphorylated AKT (17). To validate this further, we tested the effect of pharmacological blockade of AKT with two mechanically dissimilar compounds: the pan-AKT allosteric inhibitor MK2206 (AKTi^1^) and a pan-AKT ATP competitive inhibitor GDC-0068 (AKTi^2^) on proliferation of multiple *BRAF*-mutated melanoma cell lines (A375, HT144, SK-MEL28, and WM793) (23,24). Consistent with previous results, although A375, WM793 and MTG001 cells were sensitive to inhibition of all four class 1 PI3Ks with GDC0941, we observed no significant effect of AKT inhibition on melanoma cell proliferation (A375, WM793, MTG001, MTG004) regardless of PTEN status (Supp. Fig. 1). HT144 and SK-MEL28 cells were also insensitive to AKT inhibition (data not shown). In addition to the four commercially available cell lines listed above, we also present data from two cell lines derived from BRAF^V600E^-driven melanoma patient-derived xenografts (MPDX) referred to here as MTG-001 and MTG-004. These cell lines display detectable pERK and pAKT as assessed by immunoblotting (Supp. Fig. 2). Thus, pharmacological inhibition of AKT using two different compounds in a variety of cell lines has no observable effect on melanoma cell proliferation *in vitro*.

Since pharmacological inhibition was largely without effect, we attempted to silence AKT expression in *BRAF*-mutated melanoma cells using an RNA interference approach. To that end, we obtained two siRNAs targeting each of the three human AKT paralogs. A375, HT144, SK-MEL28, WM793, MTG001, and MTG004 cells were transfected with each set of siRNAs, as well as combinations thereof (siAKT1, siAKT2, siAKT3, siAKT1+2, siAKT1+3, siAKT2+3, and siAKT1+2+3) and assessed for cell confluence as a measure of proliferation using the Incucyte Live Cell Imaging System (Fig. 1A). To validate the specificity of the siRNAs, we confirmed knockdown using antibodies selective for AKT1, AKT2, and AKT3 (Fig. 1B). Silencing of individual AKT paralogs had no effect on cell proliferation when compared with a non-targeting siRNA control. Pairwise knockdown of AKT paralogs 1 and 3 resulted in a statistically significant decrease in cell proliferation in three of the cell lines tested: HT144, SK-MEL28, and MTG004 (p<0.0001) but the other pairwise combinations were without effect. However, most strikingly, siAKT1-3 resulted in complete cell lethality around 24 hours in all cell lines tested (Fig. 1A; p<0.0001). This was accompanied by significantly decreased levels of phosphorylated AKT at serine 473 (pS473) at 24 or 26 hours post transfection, and significantly decreased total AKT protein expression at 26 hours (Fig. 1C). To confirm the on-target specificity of the observed siRNA effects, we attempted a rescue experiment using mouse AKT expression vectors that are resistant to the human specific AKT1-3 siRNAs. We first validated the specificity of siAKT1-3 for human AKT in YUMM1.1 mouse melanoma cells. As expected, the siRNAs were human specific and had no effect on AKT expression in mouse melanoma cells (Supp. Fig. 3). We then transiently co-transfected the *BRAF*-mutated human cells with siAKT1-3 and vectors encoding mouse AKT1, AKT2, or AKT3. We found that expression of any of the mouse AKT paralogs could rescue the effects of the siAKT1-3 phenotype (Fig. 1C, 1D) in human melanoma cells. These data indicate that anti-proliferative effects of siAKT1-3 are on-target and any paralog of mouse AKT is able to compensate for the phenotype observed.

**Figure 1:**
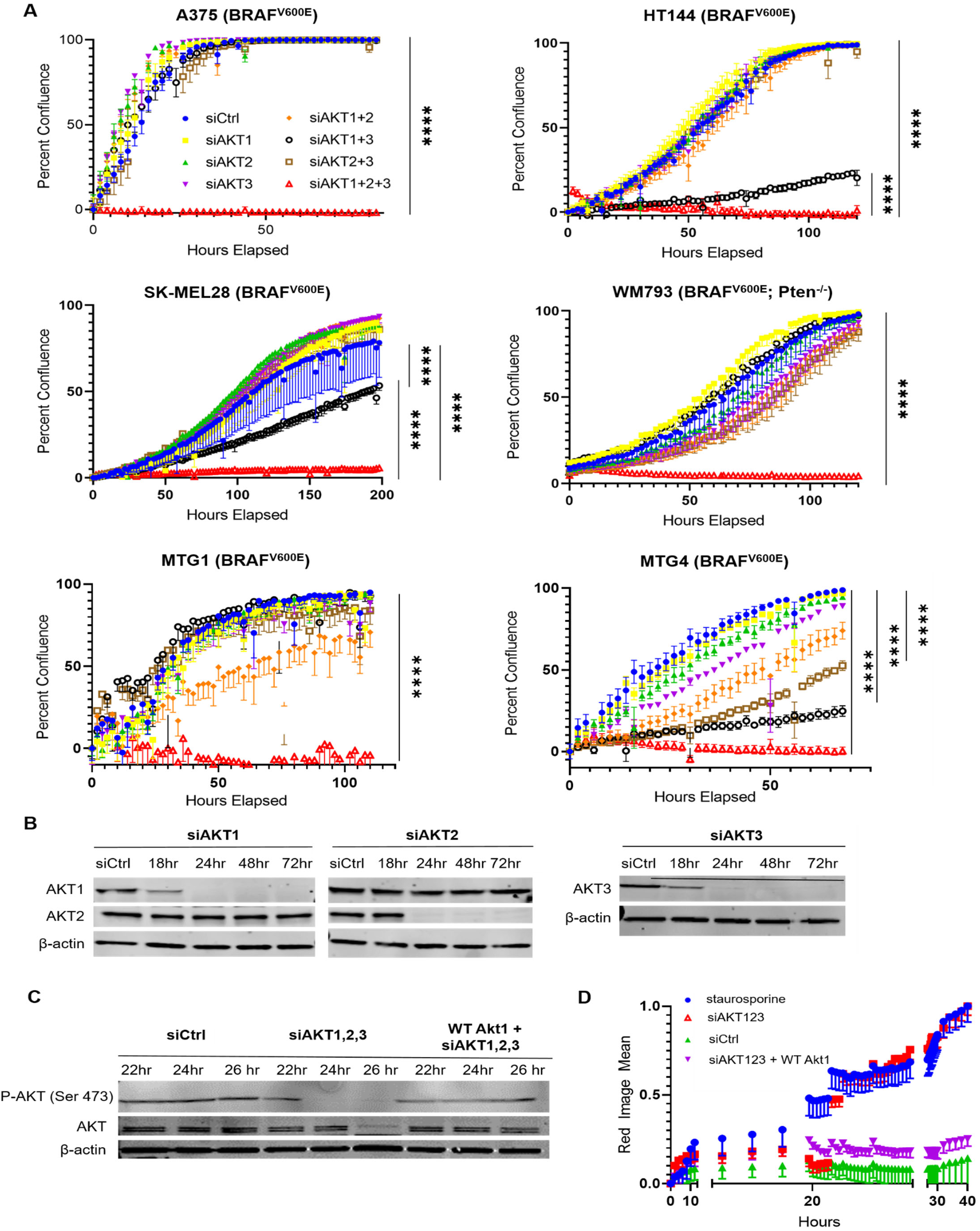
SiRNA-mediated knockdown of AKT1,2,3 leads to cell lethality. A) Cell confluency assay under genetic inhibition. A375, HT144, SK-MEL28, WM793, MTG001, and MTG004 showed little sensitivity to individual siRNAs against AKT1, 2, and 3 and AKT1+2 or AKT2+3, while siAKT123 led to complete cell lethality in all cell lines (p<0.0001 in all cell lines). HT144, SK-MEL28, and MTG004 also exhibited sensitivity to siAKT1+3 (p<0.001). Error bars indicate standard error of the mean of triplicate wells. B) Immunoblotting analysis of A375 cells treated with siAKT1 and siAKT2 in combination showed specific knockdown of individual paralogs using antibodies against total AKT1 and AKT2, as well as individual knockdown of AKT3 using siAKT3. C) Immunoblotting of A375 cells treated with siCtrl, siAKT123, and siAKT123 + overexpression of wildtype mouse Akt1 exhibited complete knockdown of phospho-AKT (Ser473) at 24 and 26 hours in siAKT123 vs siCtrl which was rescued by addition of mouse Akt1. Total AKT protein was also significantly reduced at 26 hours by siAKT123. D) Cell death assay comparing cells treated with staurosporine, siCtrl, siAKT123, and siAKT123 + WT Akt1. Treatment with siAKT123 increased cell death around 22-24 hours post transfection comparable to level of staurosporine, a multi kinase inhibitor as a control for cell death, which was completely rescued by addition of mouse WT Akt1.

To determine whether the observed decrease in cell proliferation caused by siAKT1-3 was due to induction of cell death or inhibition of cell growth, we utilized a cell death assay using CytoRed that stains permeabilized cells. Staurosporine, a pleiotropic kinase inhibitor, was used as a control for cell death. Whereas staurosporine induced a gradual increase in the percentage of dead cells over several hours, siAKT1-3 caused a dramatic increase in cell death around 22 hours, which was abrogated by ectopic expression of mouse AKT1, 2, or 3 (Fig. 1D and data not shown). Together, these data demonstrate that melanoma cell death is induced only when all three human paralogs of AKT are genetically silenced.

Recently, an AKT proteolysis-targeting chimera (PROTAC), INY-03-041, was described. This pan-AKT degrader consists of GDC-0068 conjugated to lenalidomide, a recruiter of the E3 ubiquitin ligase cereblon (25). INY-03-041 has a dual mechanism of action by both inhibiting AKT protein kinase activity (through the GDC-0068 moiety) and to target AKT proteins for proteasomal destruction (through CRBN-mediated ubiquitination); therefore, we hypothesized that it may successfully induce melanoma cell death similar to siAKT1-3. We observed ~77% reduction of total AKT expression following INY-03-041 treatment, compared with ~89% reduction by siAKT1-3, (Supp. Fig. 4A) and saw a substantial effect of 1μM INY-03-041 on melanoma cell proliferation in MTG-001 and MTG-004 cells (Supp. Fig. 4A). This effect, while not as effective as siAKT1-3, does suggest that loss of total AKT protein leads to decreased melanoma cell proliferation and may be a promising strategy for therapeutic targeting of AKT. We further hypothesize that INY-03-041 is less effective than siAKT1-3 due to less effective reduction of the individual paralogs of AKT. siRNAs targeting AKT1-3 lead to almost complete loss of each of the total paralogs (Fig. 1B). Conversely, INY-03-041 reduces AKT2 and AKT3 at 72 hours substantially, but only reduces AKT1 by 50% at 72 hours post treatment. Thus, there is less total reduction in AKT paralog expression for each of the paralogs than the individual siAKT1-3s (Supp. Fig. 4B).

### Rescue of melanoma cells from siAKT1-3 is dependent on AKT kinase activity

AKT is reported to have both kinase-dependent and kinase-independent functions. Some of its kinase-independent functions are mediated by the PH domain, which is able to sequester PI3’-lipids and influence downstream signaling (26). Having shown that mouse AKT could rescue the effects of siAKT1-3, we tested whether this rescue required AKT protein kinase activity. To that end, we generated a kinase-inactive form of mouse AKT1 in which lysine 179 was substituted with methionine to render the enzyme catalytically inactive. This mutant also harbors a deletion of the PH domain (amino acids 11-60, ΔPH) to eliminate any possible dominant-negative effect mediated by sequestration of PI3’-lipids. A myristoylation tag (amino acids 1-14 of c-SRC, myr) was added to ensure that myr-HA-ΔPH-AKT1^K179M^ was targeted to the plasma membrane for activation in the absence of a functional PH domain and an HA tag was added for immunoblot detection. A375, HT144, and WM793 cells, engineered to express myr-HA-ΔPH-AKT1^K179M^ were treated with siAKT1-3 with cell viability assessed 48 hours after siRNA transfection. In this experiment, expression of myr-HA-ΔPH-AKT1^K179M^ failed to rescue cell viability in the face of siAKT1-3, whereas a catalytically active form of this protein myr-HA-ΔPH-AKT1 demonstrated a robust rescue (Fig. 2A). This suggests that functional kinase activity, rather than kinase-independent signaling downstream of the PH domain, is required for maintaining melanoma cell viability.

**Figure 2:**
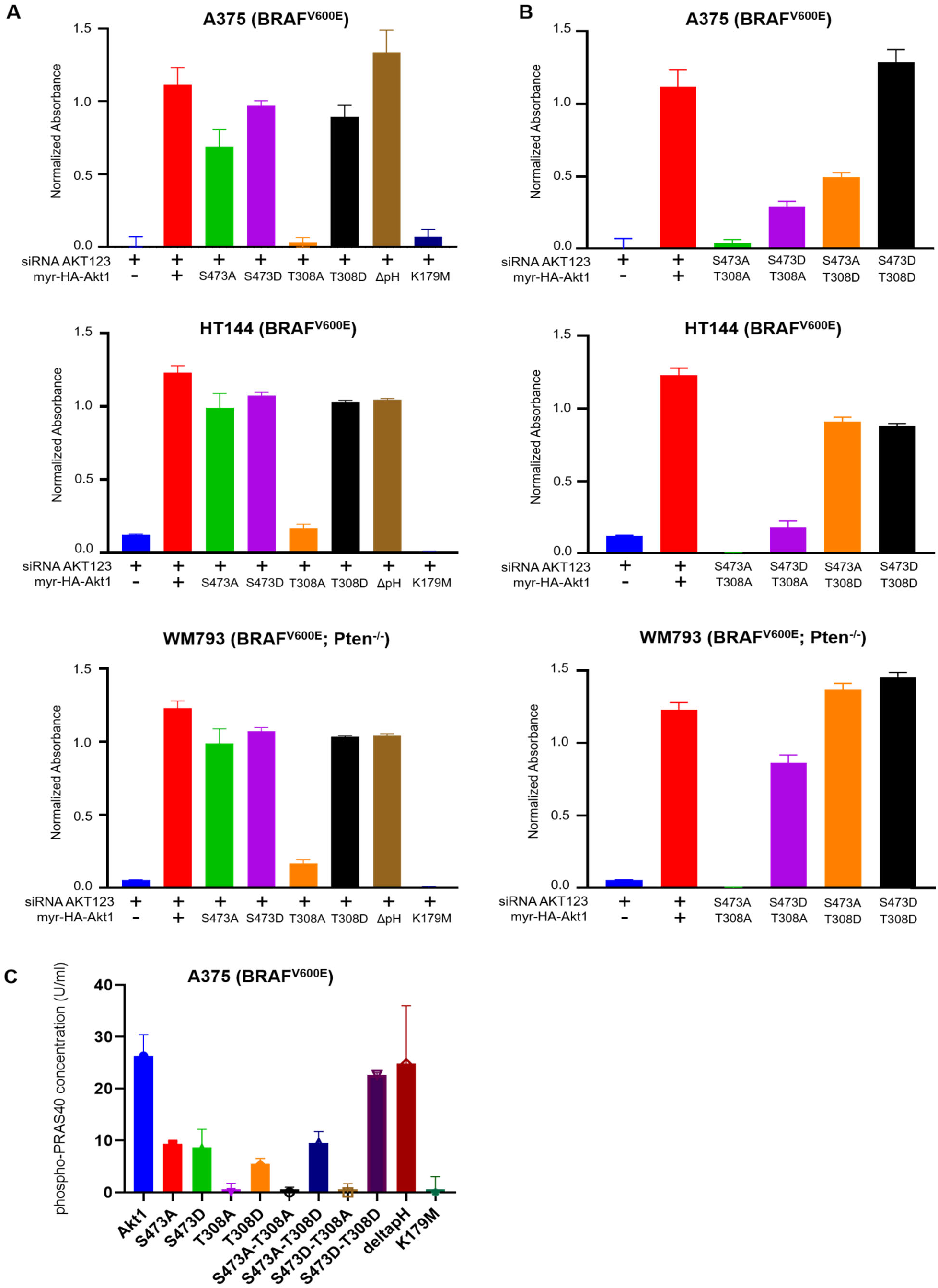
Rescue of melanoma cells from siRNA-mediated knockdown of AKT123 is dependent on Akt kinase activity and T308 phosphorylation. A) MTT assay of stable A375, HT144, and WM793 cell lines expressing phospho and kinase mutants evaluating cell viability post 48 hours siAKT123 transfection. Cell lines expressing T308A or K179M mutations were unable to significantly rescue siAKT123 knockdown while all other cell lines expressing phospho mutations were able to rescue this phenotype. B) MTT assay of stable A375, HT144, and WM793 cell lines expressing double phospho and kinase mutants evaluating cell viability post 48 hours siAKT123 transfection. C) Phospho-PRAS40 kinase ELISA of A375 myrAkt1 stable cell lines expressing phospho and kinase mutants demonstrate undetectable levels of phospho-PRAS40 in T308A; S473A, T308A; S473D, T308A; and K179M cells.

To further investigate how genetic loss of the three AKT paralogs induces melanoma cell death, we evaluated the role of AKT phosphorylation. Numerous phosphorylation sites have been identified on AKT of which three: T308, T450 and S473 are regarded as being important in the activation of AKT kinase activity. T308 is phosphorylated by PDK1, S473 by mTORC2 and T450 is thought to be a site of AKT autophosphorylation (27,28). Therefore, we investigated the role of the two major phosphorylation sites, T308 and S473, in the rescue of siAKT1-3-mediated cell death. Using site-directed mutagenesis we generated S/T>A (phospho-deficient) or S/T>D (potentially phospho-mimetic) alterations at T308 and S473 in myr-HA-ΔPH-AKT1. A375, HT144, and WM793 cells stably expressing these various forms of AKT1 were generated at which time each cell line was treated with siAKT1-3 with cell viability measured 48 hours post-transfection as described above. As expected, the two constitutively catalytically active versions of AKT, myr-HA-ΔPH-AKT1^S473D^ and myr-HA-ΔPH-AKT1^T308D^, were able to rescue cell viability. Moreover, phospho-deficient myr-HA-ΔPH-AKT1^S473A^ protein was also able to rescue cell viability. By contrast, the phospho-deficient myr-HA-ΔPH-AKT 1^T308A^ failed to rescue (Fig. 2A). These data emphasize the critical importance of T308 phosphorylation for AKT1 kinase activity

To evaluate the ability of each individual phosphomimetic and phosphodeficient mutation to cooperate with each other, we generated double point mutants: 1. myr-HA-ΔPH-AKT1^T308A/S473A^; 2. myr-HA-ΔPH-AKT1^T308A/S473D^; 3. myr-HA-ΔPH-AKT1^T308D/S473A^ and; 4. myr-HA-ΔPH-AKT1^T308D/S473D^. Perhaps as expected, the ability of these constructs to rescue the effects of siAKT1-3 was dependent on T308, as any construct with T308A failed to rescue (Fig. 2B). Interestingly, myr-HA-ΔPH-AKT1^T308D/S473A^ only achieved ~50% rescue despite the ability of individual mutants to fully rescue individually. This was a surprising but reproducible result.

To evaluate the ability of these various forms of AKT to promote phosphorylation of a direct AKT substrate, PRAS40, we used an ELISA assay to assess pT246-PRAS40 in suitably prepared cell extracts. As expected, pT246-PRAS40 was largely undetectable in cells expressing myr-HA-ΔPH-AKT1^K179M^, myr-HA-ΔPH-AKT1^T308A/S473D^ or myr-HA-ΔPH-AKT1^T308A/S473A^ in accordance with the inability of these mutants to rescue siAKT1-3 (Fig. 2C). By contrast, pT246-PRAS40 was readily detected in those cells in which the various AKT1 constructs successfully rescued the effects of siAKT1-3: myr-HA-ΔPH-AKT1, myr-HA-ΔPH-AKT1^T308D/S473A^ and myr-HA-ΔPH-AKT1^T308D/S473D^.

### siAKT1-3 significantly decreases mTORC activity

To identify downstream signaling effectors that lead to cell death upon treatment with siAKT1-3, we utilized the Luminex® xMAP® quantitative immunoassay technology to measure phosphorylation of a panel of cell signaling molecules in the PI3K>AKT>mTORC signaling pathway. Measurement of pS473-AKT was used as a marker of catalytically activated AKT1-3 since this antibody recognizes all three AKTs when phosphorylated on the cognate sites S474-AKT2 and S472-AKT3). Consistent with published literature, treatment of MTG-001 and MTG-004 cells with inhibitors of PI3’-kinase (GDC0941 or BYL719), or of AKT (MK-2206 or siAKT1-3) led to decreased pS473-AKT at 22 or 24 hours after treatment (Fig. 3A). Treatment of cells with GDC-0068 or INY-03-041 increased phosphorylated pS473-AKT consistent with their proposed mechanism of hyperphosphorylation leading to a locked, inactive conformation. In cells treated with siAKT1-3, we noted substantially decreased levels of phosphorylation of proteins downstream of mTORC1, including pT389-p70^S6K^, pS235-RPS6 and pS2448-mTORC in cell lines derived from the MTG001 or MTG004 MPDX models. Interestingly, compared with the AKT inhibitors MK2206 and GDC0068, siAKT1-3 at 24 hours substantially reduced pT389-p70^S6K^ (Fig. 3B, p<0.0001 in both MTG001, MTG004), pS235-RPS6 (Fig. 3C), and pS2448-mTORC (Fig. 3D). These results were verified by immunoblotting and compared with their respective total levels of each protein (Supp. Fig. 4B). In addition to noting consistent differences in pS473-AKT and pS235-RPS6, we also noted decreased levels of phosphorylated PRAS40 and total AKT in siAKT1-3 at 24 hours compared with the pharmacological inhibitors in each of the MTG001 and MTG004 cell lines. In addition, siAKT1-3 displayed highly diminished levels of 4EBP1 (pThr37/46) downstream of mTORC1. These data suggest that AKT acts to promote cell viability through direct mTORC1-dependent mechanisms.

**Figure 3:**
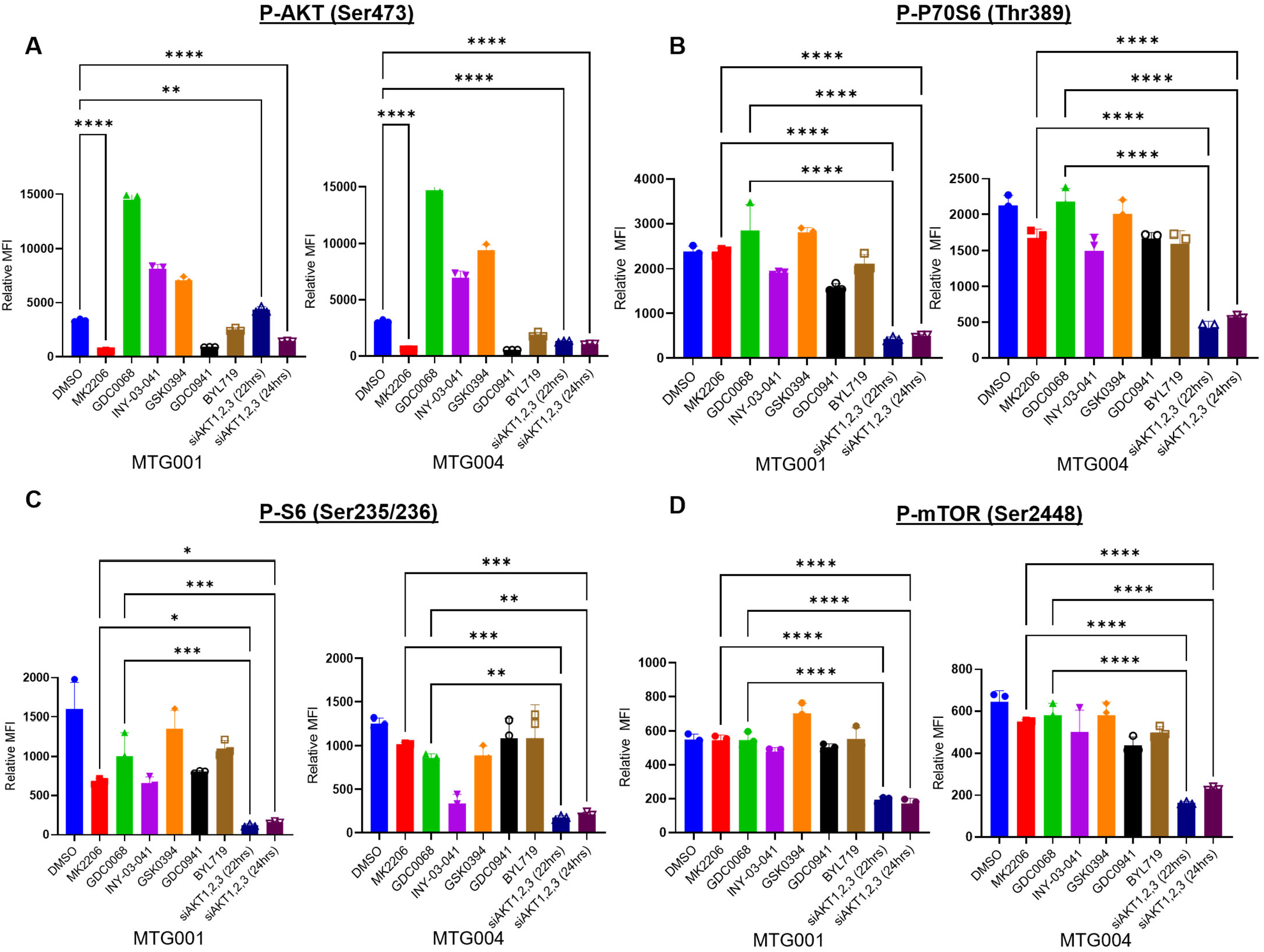
SiAKT123 significantly decreases mTORC activity. A) A-D, Luminex quantitative immunoassay of phosphorylation of AKT (Ser473), P70S6 (Thr389), P-S6 (Ser235/236A, MTG001 and MTG004 treated with DMSO control, MK2206 (2.5μM), GDC0068 (1μM), INY-03-041 (500nM), GDC0941 (1μM), BYL719 (1μM), siAKT123 at 22 hours (50nM), or siAKT123 at 24 hours (50nM). MK2206, GDC0941, and siAKT123 (24 hours) significantly decreased p-AKT (p<0.0001 in both cell lines) as expected compared to DMSO control. SiAKT123 (22 hours) significantly decreased pAKT compared to DMSO control in MTG004 (p<0.0001). GDC0068, INY-03-041, and GSK0394 significantly increased pAKT compared to DMSO control (p<0.0001 for all treatments). B) SiAKT123 at 22 and 24 hours significantly decreased p-P70S6 compared to MK2206 and GDC0068 in both MTG001 and MTG004 (p<0.0001). C) SiAKT123 (22 hours) and siAKT123 (24 hours) decreased p-pS6 in comparison to MK2206 (p=0.02 and p=0.04 respectively) and in comparison to GDC0068 in MTG001 (0.0002 and 0.0004 respectively). SiAKT123 (22 hours) and siAKT123 (24 hours) decreased p-pS6 in comparison to MK2206 (p=0.0001 and 0.0002 respectively) and in comparison to GDC0068 (0.0011 and 0.0020 respectively) in MTG004. D) SiAKT123 at 22 and 24 hours significantly decreased p-P70S6 compared to MK2206 and GDC0068 in both MTG001 and MTG004 (p<0.0001).

### Exogenous expression of SGK rescues the lethal effect of siAKT1-3

It has previously been shown that SGK signals downstream and is activated by the PI3K signaling cascade and also that, like AKT protein kinases, mTORC1 directly phosphorylates SGK1 leading to its activation (29). Hence, we investigated whether ectopic expression of either normal or constitutively active, myristolated SGK could rescue siAKT1-3 in melanoma cells. Compared with our negative control, stable melanoma cell lines overexpressing either mouse AKT1, or myristolated forms of SGK1, 2, or 3 were all able to significantly rescue the lethal effect of siAKT1-3 (Fig. 4A, B, p<0.0001). Ectopic expression of SGK plus the effects of siAKT1-3 were verified both by qRT-PCR and immunoblotting (Fig. 4C and D). Compared with siAKT1-3-treated alone, normal SGK1 was unable to rescue this phenotype suggesting that constitutive activation of the SGK protein kinase is required to rescue the lethal effects of siAKT1-3 (Fig. 4E, p=0.8758).

**Figure 4:**
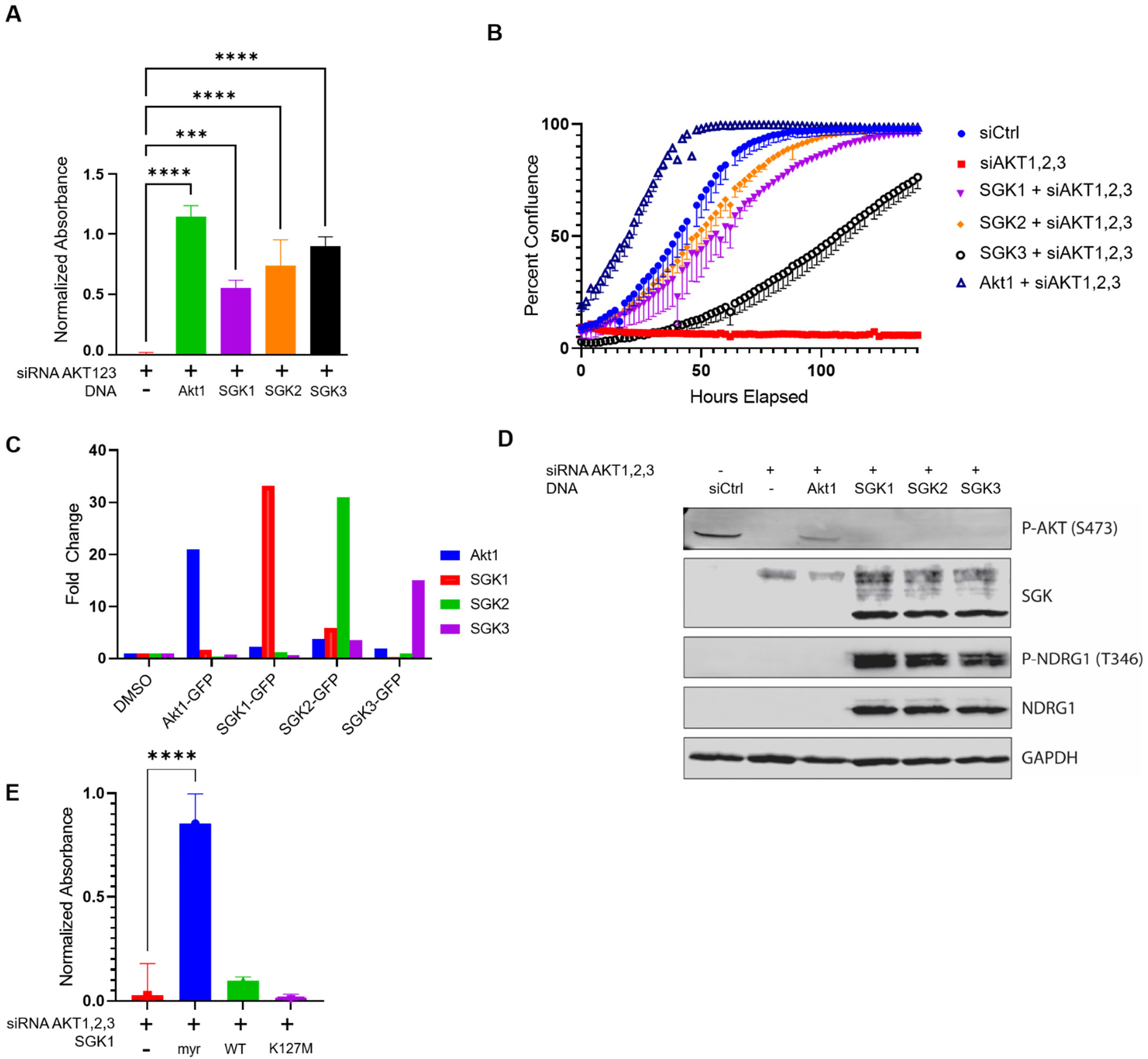
SGK can rescue the lethal effect of siRNA-mediated knockdown of AKT1,2,3. A) MTT assay of stable A375 cell lines expressing myrAkt1, myrSGK1, myrSGK2, or myrSGK3 evaluating cell viability post 48 hours siAKT123 transfection compared to siAKT123 alone. MyrAkt1, myrSGK1, myrSGK2, and myrSGK3 were all able to significantly rescue cell death (p<0.0001). B) Cell proliferation assay evaluating cell proliferation in A375, A375 myrAkt1, A375 myrSGK1, A375 myrSGK2, and A375 myrSGK3 cell lines treated with siCtrl or siAKT123. A375 myrAkt1, A375 myrSGK1, A375 myrSGK2, and A375 myrSGK3 rescued cell proliferation in presence of siAKT123 knockdown. C) Q RT-PCR demonstrating increased specific gene expression of Akt1, SGK1, SGK2, and SGK3 in A375 stable cell lines. D) Immunoblotting of A375, A375 myrAkt1, A375 myrSGK1, A375 myrSGK2, and A375 myrSGK3 treated with siCtrl or siAKT123 demonstrating knockdown of p-AKT except for in A375 myrAkt1 cell line. SGK and p-NDRG1 expression is increased in A375 myrSGK1, A375 myrSGK2, and A375 myrSGK3 cells. E) MTT assay of A375 myrSGK1, A375 wt SGK1, and A375 myrSGK1-K127M cell lines evaluating cell viability post 48 hours siAKT123 transfection. Only A375 myrSGK1 could rescue siAKT123 knockdown.

As we found that the kinase activity of mouse AKT1 was indispensable for its ability to rescue siAKT1-3 effects, stable melanoma cell lines with ectopic expression of SGK1 containing catalytically inactive mutations (K127M and K191M respectively) were generated to evaluate the necessity of SGK kinase activity to rescue siAKT1-3 knockdown. Compared with ectopic expression of constitutively active SGK1, a kinase-dead version was unable to rescue the lethal effects siAKT1-3 (Fig. 4E). Together, these results suggest that the protein kinase activity of SGK is able to compensate for the silencing of all AKT protein kinases in melanoma cells.

To further assess the role of SGK in this context, we obtained siRNAs targeting each of the SGK paralogs and tested the consequences of genetic silencing of SGK1-3 to inhibit melanoma cell viability. Interestingly, siRNAs targeting SGK1, 2, or 3, or any pairwise or the triple combination, had no effect on melanoma cell proliferation suggesting that while SGK is able to rescue the effects of siAKT1-3 (Supp. Fig. 5), AKT is the predominant activator of the mTORC signaling that is necessary for melanoma cell viability.

### SGK1 enhances melanoma tumor growth *in vivo*

RNA sequencing analysis of matched nevi vs primary melanoma samples revealed that SGK1 mRNA is significantly elevated in melanoma (p=0.0001) (Supp. Fig. 6). Moreover, analysis of RNA or proteins from siAKT1-3 treated MTG-001 and MTG-004 cells at 22 hours demonstrated a significant increase in SGK1 mRNA expression and total SGK1 protein levels, as well as increased phosphorylation of NDRG1 (p-Thr346), a known downstream target of SGK1. Therefore, we investigated the role of SGK1 in promoting melanoma progression *in vivo*. To that end, we utilized the YUMM3.2 mouse melanoma cell line (<u>B</u>RAF^V600E^/<u>I</u>NK4A-<u>A</u>RF^Δ^), as well as a PTEN-deficient isogenic variant (BRAF^V600E^/INK4A-ARF^Δ^/PTEN^Δ^) of this line generated using CRISPR/CAS9 technology. These isogenic variants were then used to generate stable cell lines ectopically expressing myristoylated SGK1 along with luciferase and then injected into immune-competent syngeneic mice for *in* vivo analysis.

We observed that ectopic expression of SGK1 was able to potentiate melanoma growth regardless of the status of PTEN. Specifically, ectopic expression of SGK1 in cells either proficient or deficient in expression of PTEN decreased time to tumor onset and reduced overall survival. Tumors with functional PTEN had an average overall survival of 32.5 days with a median survival of 32 days. By comparison, tumors with functional PTEN and ectopic expression of SGK1 significantly reduced median and average overall survival compared to the parental cell line (Fig. 5A, p<0.001). Additionally, ectopic expression of SGK1 led to reduced tumor latency (Fig. 5B, 5C) such that tumors with loss of PTEN had an overall survival of 30.6 days and a median survival of 28 days compared to those with ectopic expression of SGK1 (22.9 days and 24.5 days respectively). Thus, SGK1 combined with loss of PTEN significantly reduced overall and median survival (Fig. 5D, p = 0.03) and led to reduced time to tumor onset (Fig. 5E, 5F). These data demonstrate that SGK1 is able to potentiate melanoma tumor growth and decrease overall survival regardless of PTEN status.

### Combined inhibition of AKT and SGK decreases melanoma cell proliferation

Since the effects of siAKT1-3 were rescued by ectopic expression of SGK1, we evaluated whether combined pharmacological inhibition of AKT and SGK would decrease melanoma cell proliferation. To that end, we employed GSK650394, a pharmacological inhibitor of SGK that selectively inhibits SGK1 and 2 but has very limited reported inhibitory effects on AKT (30). MTG001 and MTG004 melanoma cells were treated with: 1. Vehicle control; 2. MK2206 (2.5μM, AKTi^1^), GDC-0068 (1μM, AKTi^2^), INY-03-041 (500nM, AKT PROTAC), GSK650394 (3μM, SGK inhibitor), or the combination of GSK650394 plus each of the various AKT inhibitors. As expected, treatment with the AKT inhibitors did not lead to a significant decrease in cell proliferation. By contrast, treatment with the SGK inhibitor GSK650394 alone yielded a statistically significant decrease in cell proliferation. However, the cells continued to grow, albeit more slowly, throughout the treatment. Strikingly, treatment with the combination of the SGK inhibitor together with either of the AKT inhibitors or PROTAC all led to cell stasis and the absence of continued cellular proliferation (Fig. 6A). Thus, while AKT inhibitors alone are ineffective, their activity is enhanced when used in combination with SGK inhibitors to block melanoma cell proliferation.

**Figure 5:** SGK1 enhances melanoma tumor growth *in vivo*. A) Immune competent mice tolerized to luciferase and GFP were injected subcutaneously with YUMM 3.2 mouse melanoma cells (parental or myrSGK1). There was a significant decrease in survival for mice whose primary tumors overexpressed myrSGK1 (red) compared with empty vector controls (blue) (p<0.0001). B) Mice were injected with YUMM 3.2 Pten deficient mouse melanoma cells (parental or myrSGK1). There was a significant decrease in overall survival for mice whose primary tumors overexpressed myrSGK1 (red) compared to empty vector controls (blue) (p=0.03). A logrank (Mantel-Cox) test was used to determine significant differences in survival. C) Tumor growth curves for mice injected with YUMM 3.2 myrSGK1 demonstrated decreased tumor latency compared to YUMM 3.2 empty vector control. D) Tumor growth curves for mice injected with YUMM 3.2 Pten null myrSGK1 also demonstrated decreased tumor latency compared to YUMM 3.2 empty vector control. E) Mice were injected with YUMM 3.2 parental or YUMM 3.2 myrSGK1 cells that express luciferase and were imaged at one-week intervals via BLI. Mice that expressed myrSGK1 primary tumors had higher luciferase activity at earlier weekly timepoints compared to empty vector controls. By Week 3, myrSGK1 mice had to be sac’d due to tumor burden in Pten wildtype cohorts. F) Mice that were injected with YUMM 3.2 Pten null myrSGK1 had higher luciferase activity at earlier timepoints compared to YUMM 3.2 Pten null empty vector.

**Figure 6:**
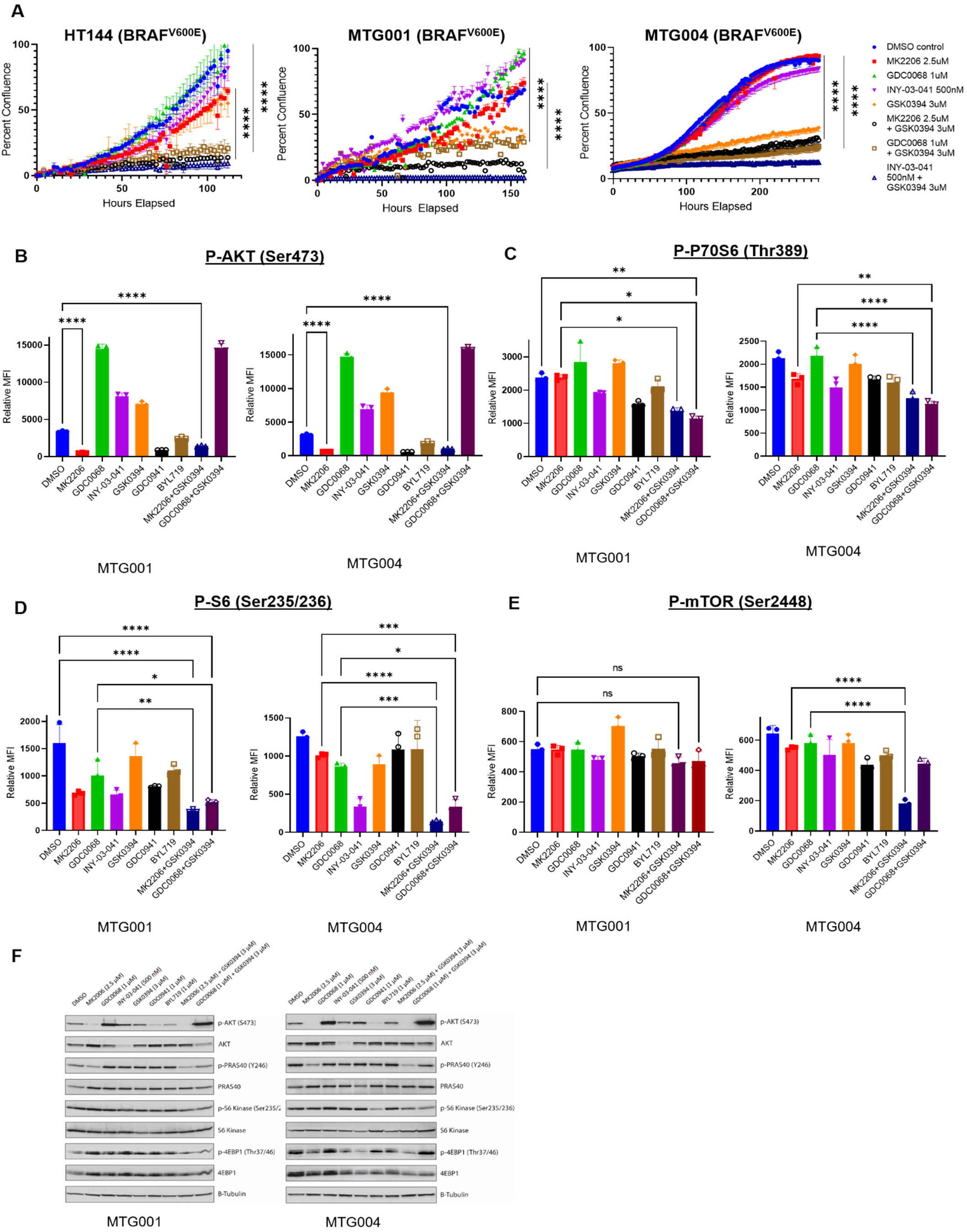
Combined inhibition of AKT and SGK decreases melanoma cell proliferation through effects on mTORC signaling. A) Cell confluency assay under pharmacological inhibition. HT144, MT001, and MTG004 cells were treated with DMSO control, MK2206 (2.5μM), GDC0068 (1μM), INY-03-041 (500nM), GSK0394 (3μM), MK2206 (2.5μM) + GSK0394 (3μM), GDC0068 (1μM) + GSK0394 (3μM), and INY-03-041 (500nM) + GSK0394 (3μM). Treatment with the combination treatments of MK2206 + GSK0394, GDC0068 + GSK0394, and INY-03-041 + GSK0394 led to significantly decreased cell proliferation in all three lines (p<0.0001). Treatment with GSK0394 alone also led to statistically decreased cell proliferation in MTG001 and MTG004 (p<0.0001). B) B-E, Luminex quantitative immunoassay of phosphorylation of AKT (Ser473), P70S6 (Thr389), P-S6 (Ser235/236A, MTG001 and MTG004 treated with DMSO control, MK2206 (2.5μM), GDC0068 (1μM), INY-03-041 (500nM), GDC0941 (1μM), BYL719 (1μM), MK2206 (2.5μM) + GSK0394 (3μM), or GDC0068 (1μM) + GSK0394 (3μM). B, MK2206, GDC0941, and MK2206 + GSK0394 significantly decreased pAKT (p<0.0001), while GDC0068, GSK0394, INY-03-041, and GDC0068 + GSK0394 all significantly increased pAKT (p<0.0001) in both MTG001 and MTG004. C) Double combination of MK2206 + GSK0394 or GDC0068 + GSK0394 significantly decreased p-p70S6 compared to MK2206 (p=0.042 and p=0.0124) in MTG001. GDC0068 + GSK0394 significantly decreased p-p70S6 versus either single AKT inhibitor (MK2206 or GDC068) (p=<0.0001) in MTG004. D) Both combination treatments (MK2206 + GSK0394 or GDC0068 + GSK0394) decreased p-S6 compared to GDC0068 alone in MTG001 (p=0.0039 and p=0.0468 respectively). Combination treatments decreased p-S6 compared to both single AKT agent alone in MTG004 (vs MK2206: p<0.0001 and p=0.0008, vs. GDC0068: 0.0004 and 0.0110). E) There was no significance in decrease of mTOR phosphorylation by combination treatments versus single agent inhibition in MTG001, but MK2206 + GSK0394 resulted in significant decrease in p-mTOR versus either single AKT agent alone in MTG004 (p<0.0001). F) Immunoblotting analysis of lysates treated with pharmacological inhibition and used for Luminex analysis (B-E). Decreased levels of p-PRAS40, p-S6, and p-4EBP1 with combination treatments versus single agent alone were observed, particularly in MTG001.

To investigate the downstream mechanisms involved in decreased melanoma cell proliferation with the combined pharmacological treatments, we again utilized the Luminex® xMAP® quantitative immunoassay technology to measure phosphorylation of a panel of eight cell signaling molecules in the PI3K>AKT>mTORC signaling pathway as described above. Again, pS473-AKT confirmed the mechanism of action of each pharmacological inhibitor treatment. Interestingly, treatment with GSK650394 yielded a statistically significant increase of pS473-AKT (Fig. 6B), just as inhibition of AKT via siAKT1-3 yields an increase of SGK1 expression suggesting further compensatory feedback between these two signaling proteins. Importantly, downstream phosphorylation of pT389-p70^S6K^ (Fig. 6C) and pS235/236-RPS6 (Fig. 6D) were significantly decreased in each of the MTG-001 or MTG-004 lines with the combined pharmacological treatment of MK2206 plus GSK650394, as well as GDC-0068 and GSK650394. MK2206 and GSK650394 yielded a statistically significant decrease of p-mTOR at Ser2448 compared to both of the single agent AKT inhibitors, MK2206, and GDC-0068, alone in MTG004 (Fig. 6E). These lysates were further evaluated by immunoblotting and compared to total levels of protein, showing evidence of lower levels of p-PRAS40 (Y248), p-S6 (Ser235/236), and p-4EBP1 (Thr37/46) with combination treatments versus single agent alone, particularly in MTG-001 (Fig. 6F). Based on this data, we conclude that single agent pharmacological inhibition of AKT does not lead to effective suppression of downstream mTORC signaling necessary to inhibit melanoma cell proliferation. However, combined suppression of both AKT and SGK led to decreased melanoma cell growth.

## DISCUSSION

The striking success of inhibitors of BRAF^V600E^>MEK>ERK signaling in patients with *BRAF*-mutated melanoma begs the question as to the utility of additional pathway-targeted agents in this disease, either as single agents or in combination therapy, in which patients might be stratified using predictive biomarkers. Examples include the possible use in melanoma of: CDK4/6 inhibitors in tumors with INK4A silencing; MDM2 inhibitors in tumors that express normal TP53 or; PI3K or AKT inhibitors in tumors with alterations in NRAS, RAC1, PIK3CA, AKT1-3 or PTEN (31). However to date, despite compelling preclinical data supporting their clinical deployment, the use of PI3K or AKT inhibitors in the treatment of melanoma has been confounded by issues of toxicity, lack of efficacy or both (32,33). Furthermore, although there are five PI3’-kinase inhibitors FDA approved for cancer or PI3Kα-related overgrowth spectrum (alpe-, copan-, duve-, idela- and umbralisib), there are no AKT inhibitors FDA approved for cancer therapy, albeit that GDC-0068 (ipatasertib) is showing promising activity in prostate cancer clinical trials (34,35). Hence, we sought to elucidate the role of AKT in melanoma cell proliferation. However, our results are somewhat paradoxical in that, although pharmacological inhibition of AKT1-3 protein kinase activity had little or no effect on melanoma proliferation, rescue of the potent anti-proliferative activity of siAKT1-3 was apparently dependent upon the catalytic activity of mouse AKT. Moreover, the fact that the expression of all three AKT paralogs had to be silenced for the effects of siAKT1-3 suggests a substantial degree of overlap in AKT function in melanoma cell proliferation. SiRNAs targeting AKT1,2&3 resulted in loss of cell viability, which was rescued by ectopic expression of mouse AKT, regardless of paralog. This was dependent on functional kinase activity and phosphorylation of T308. Several of our previous studies focused on better understanding the independent non-overlapping functions of each AKT paralog in melanoma initiation and progression. For example, we previously reported that a major hotspot mutation in AKT1, E17K, leads to melanoma metastasis (9). However, here we show that each individual paralog (AKT 1, 2, or 3) has overlapping functions that each contribute to melanoma cell growth and survival.

Although AKT activation has been shown to require phosphorylation of both T308 and Ser473, multiple sources have debated the necessity of phosphorylation at each of these two residues for full activation (27). Key pioneering research by Alessi et al. reported that both phosphorylation by PDK1 and mTORC2 is necessary for full activation (36,37). In our context, based on the readout of siAKT1-3 rescue, phosphorylation of S473 appears largely dispensable for the ability of activated AKT to rescue the lethal effects of siAKT1-3.

Prior research has suggested a role for SGK protein kinases in melanoma, but we report the first evidence that activated SGK1 can promote melanoma tumorigenesis. Previous studies demonstrated that SGK is a key mediator of PDK1 activity in melanoma *in vitro* studies, particularly in PTEN proficient melanoma cells, in which there were low levels of phospho-AKT (38). To that end, we tested the ability of constitutively activated (CA) SGK1 to promote tumorigenesis in isogenic YUMM3.2 (BRAF^V600E^/INK4A-ARF^Δ^) cells that were either proficient or deficient for PTEN expression. Interestingly, CA-SGK1 could promote tumorigenic growth of YUMM3.2 cells regardless of PTEN status leading more rapid onset of end-stage disease and a decrease in overall survival. However, although these data highlight the pro-tumorigenic capability of SGK1, it is not clear if SGK1 plays a role in tumorigenic growth of BRAF-mutated melanomas in humans. However, although not mutated in melanoma per se, SGK1 mRNA expression is noted to increase in the context of the transition of nevi to melanoma.

SGK1 expression is increased in the context of siAKT1-3 knockdown, but it is evident that endogenous wild type SGK1 is not sufficient to rescue abrogation of cell proliferation by siAKT1-3. We hypothesize that wild type SGK requires mTORC1 for activation, and this signal is absent in cells treated with siAKT1-3. Constitutively active SGK is independent of upstream mTORC1 activation and is therefore able to rescue the phenotype in this context.

Activation of mTORC signaling complexes most likely occurs in a majority of malignant melanomas since most melanomas display phosphorylation of direct mTORC targets such as 4E-BP1 and p70^S6K^ (39). Indeed, evidence from this study shows that *BRAF*-mutated melanomas, regardless of PTEN status, rely on mTORC1 signaling. However, mTORC activation is often under the dual control of both the BRAF^V600E^>MEK>ERK and the PI3’-kinase signaling pathways (40).Consistent with this hypothesis, we report here that single agent pharmacological inhibition of AKT or SGK is insufficient to fully diminish mTORC signaling. In other cancer lines, studies have also suggested that activity of either AKT or SGK1 is sufficient to mediate signaling through mTOR (41), which we also observed in melanoma. AKT and SGK both phosphorylate p27 thus blocking nuclear import and maintaining active cyclin-E/Cdk2 complexes (29,42). However, only SGK is directly phosphorylated by mTORC1. Our results propose that in the absence of AKT due to genetic targeting, mTORC1 activity is extinguished, preventing phosphorylation of SGK and subsequently p27. Based on our results, we further suggest that AKT is the predominant kinase whose cell proliferation activity through phosphorylation of p27 must be fully extinguished to lead to complete suppression of mTOR as our data demonstrates that genetic suppression of SGK123 does not have the same effect on cell viability.

Despite our results suggesting that key proliferation of melanoma cells is through effects on mTOR, phase II clinical trials of mTOR inhibitors have not shown clinical advantage. This may be due to multiple reasons: firstly, mTOR inhibitors, such as rapamycin, function by destabilization of the mTORC1-Raptor complex while leaving the mTORC2-Rictor complex, intact. Rictor enables mTORC2 to directly phosphorylate pS473-AKT and facilitates T308 phosphorylation by PDK1 (29). As both AKT and SGK are phosphorylated by mTORC2 and PDK1 to facilitate downstream signaling through mTORC1, residual activity of mTOR incompletely suppressed by rapamycin may still be sufficient to drive melanoma progression. With the current inadequacy of PI3K pathway inhibitors, including AKT inhibitors, to effectively target all downstream nodes of the pathway, notably mTOR, our study suggests that, if tolerable, combined inhibition of AKT plus SGK protein kinases may represent a useful combination treatment strategy to more effectively target these signaling pathways and eliminate upstream compensatory signaling networks that may lead to resistance.

## ACKNOWLEDGEMENTS

We thank members of the VanBrocklin, Kinsey, McMahon, and Holmen labs as well as A. Welm and R. Stewart for providing mouse strains, reagents, vectors and/or advice. We thank Nathaniel Gray and Alex Toker for their generous gift of the INY-03-041. We thank HCI Shared Resources (including Flow Cytometry, Histology, DNA sequencing) for their support and help. GP, MM, and SH were supported by grants from NIH (F31CA254307, CA121118, and CA176839) and institutional funds (Huntsman Cancer Foundation). GP was supported by a JEDI award from the Life Sciences Editors Foundations, and we graciously thank Helen Pickersgill for editing suggestions and advice.

## AUTHOR CONTRIBUTIONS

GP, SH, and MM designed experiments. GP, TT, DK, WB, CM, KO, RF, RE, and MF performed experiments. GP, DK, WB, CM, SH, and MM analyzed data. GP, SH, and MM wrote the manuscript. All authors discussed results, reviewed and revised the manuscript.

---

**Supplemental Figure 1:**
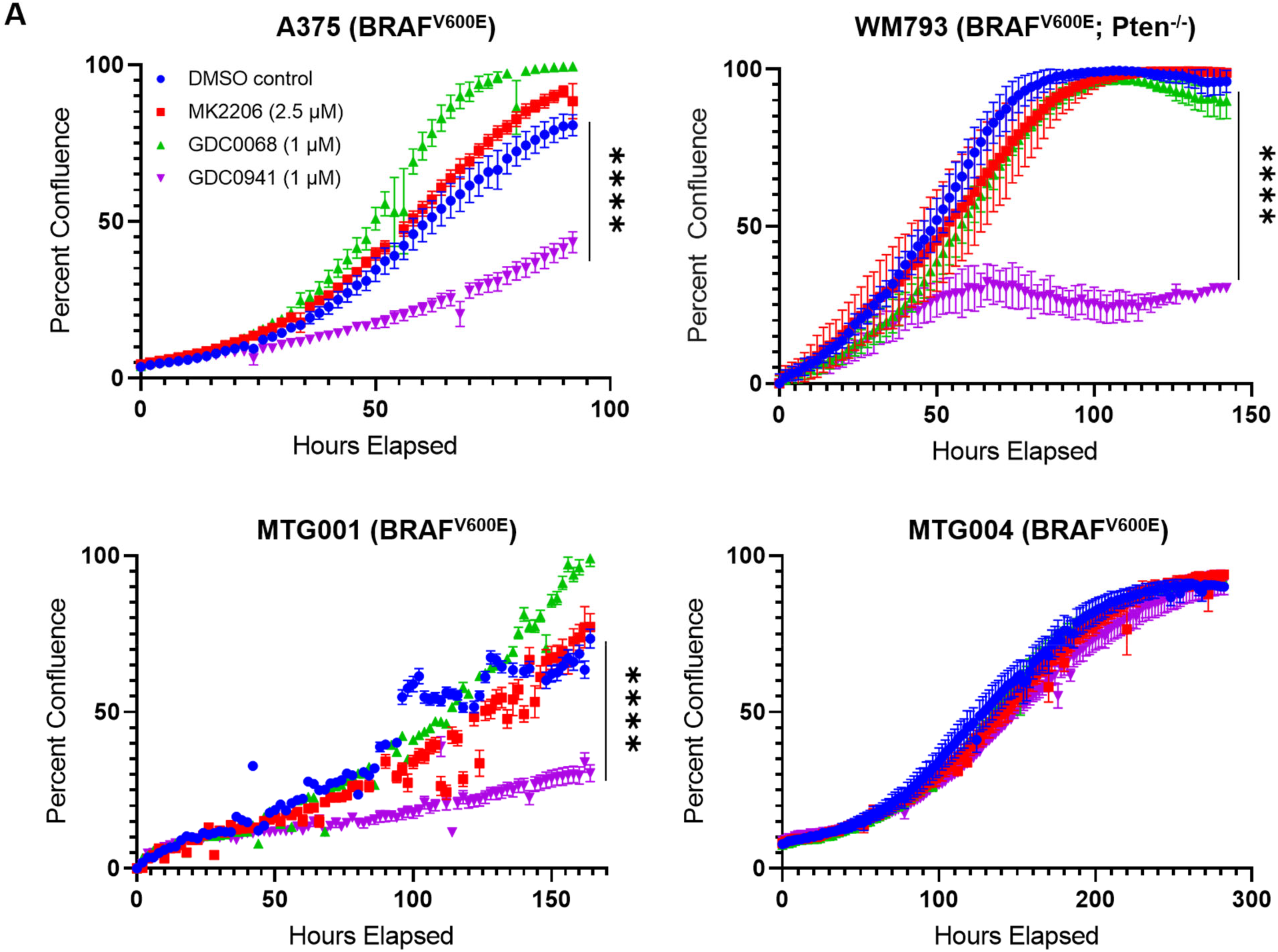
Pharmacological inhibition has little effect on melanoma cells. A) Cell confluency assay under pharmacological inhibition. A375, WM793, MTG001, and MTG004 cells were largely resistant to AKT inhibitors MK2206 (2.5 μM) and GDC0068 (1μM) while A375, WM793, and MTG001 cells were largely sensitive to PI3Kα inhibition by GDC0941 (1μM) (p<0.001). Error bars indicate standard error of the mean of triplicate wells.

**Supplemental Figure 2:**
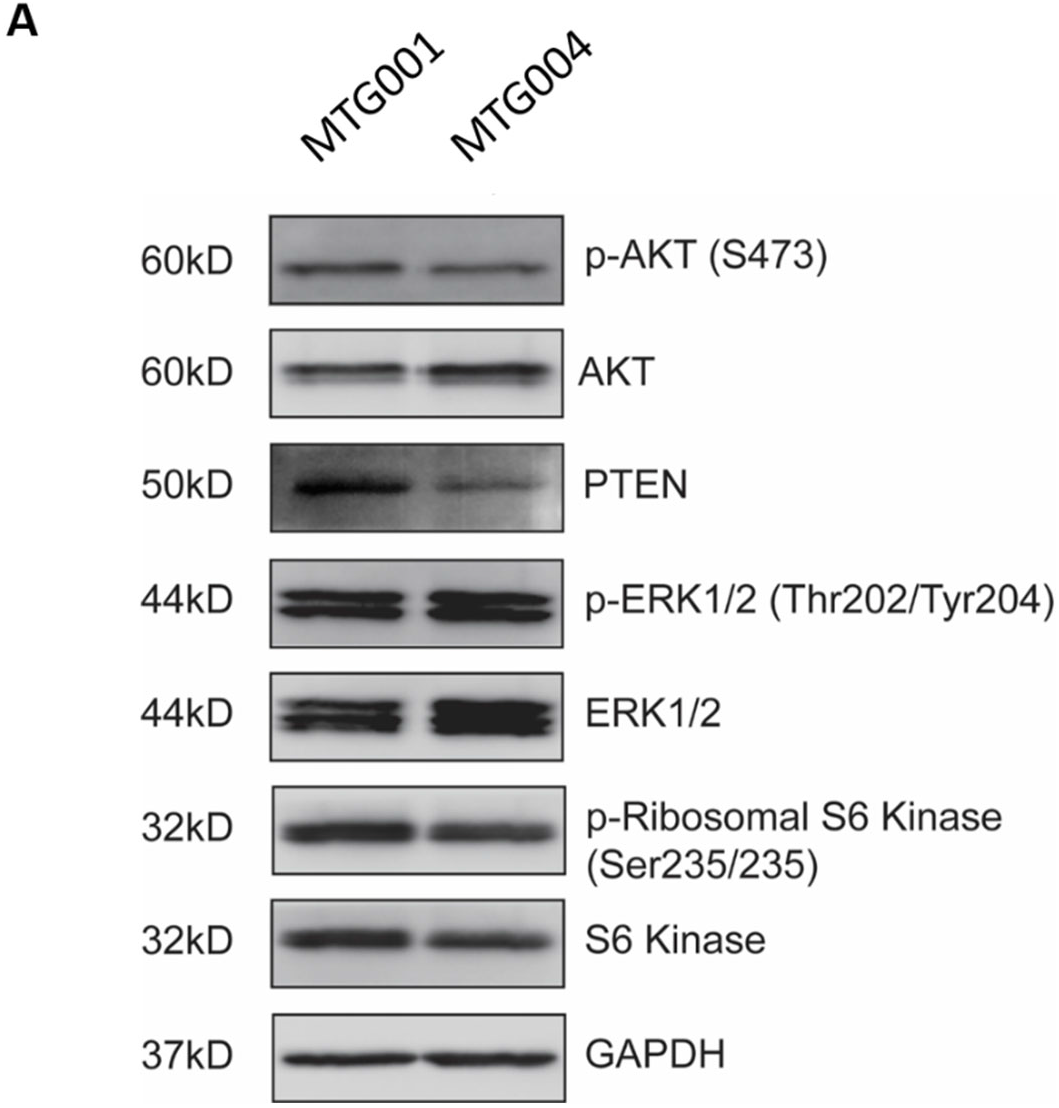
Classification of MTG001 and MTG004. A) Immunoblotting of MTG001 and MTG004, PDX-derived cell lines, exhibited similar levels of phospho-AKT (Ser473), total AKT, pERK (Thr202/Tyr204), ERK1/2, p-Ribosomal S6 Kinase (Ser235/236), S6 Kinase. GAPDH was used as a loading control. MTG001 exhibits higher levels of total PTEN.

**Supplemental Figure 3:**
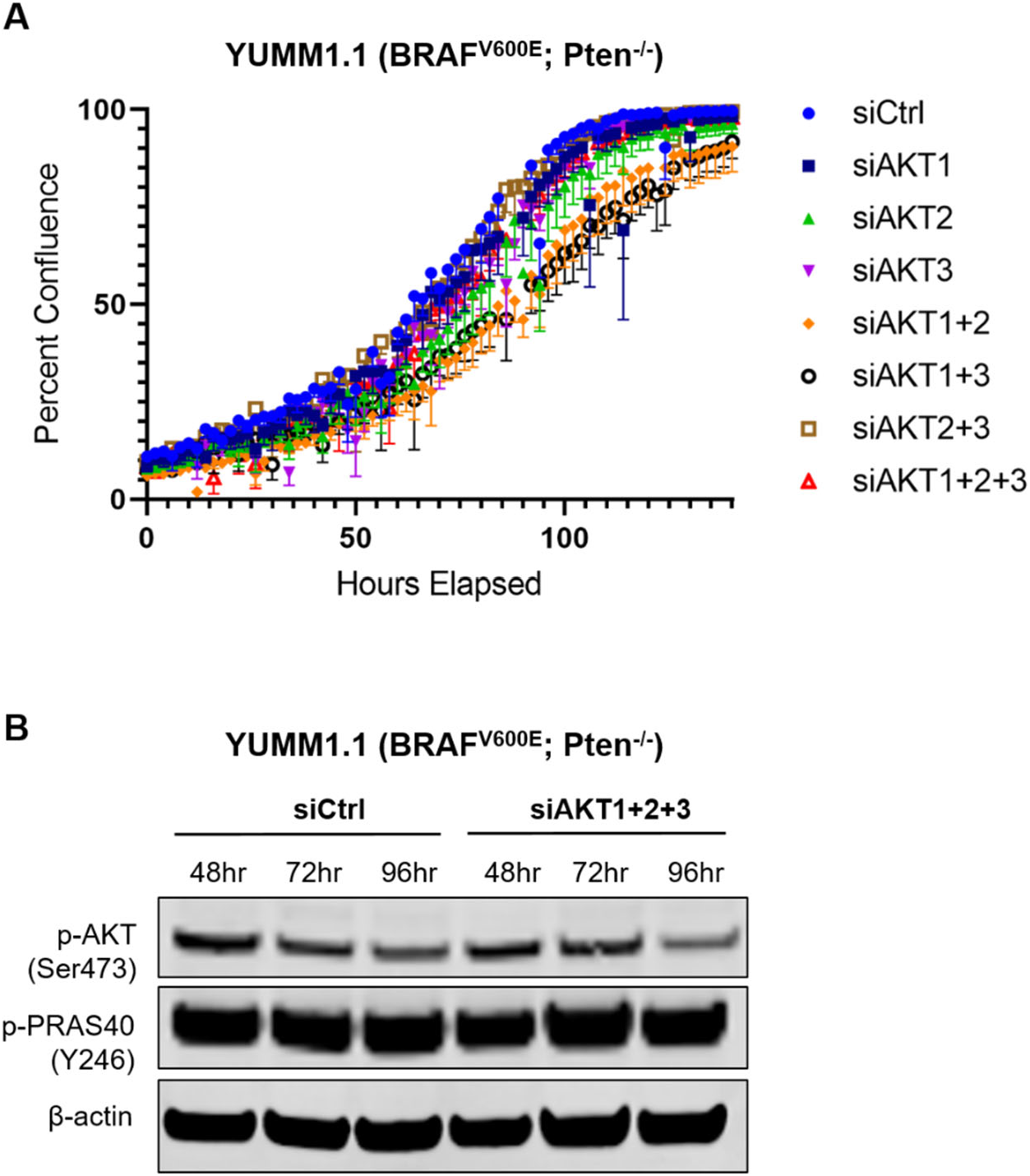
SiAKT123 is specific for human AKT. A) Cell confluency assay of YUMM1.1 mouse melanoma cells under genetic inhibition demonstrating no effect on cell death with siAKT1, siAKT2, siAKT3, or any combination. Error bars indicate standard error of the mean of triplicate wells. B) Immunoblotting of YUMM1.1 cells treated with siCtrl vs siAKT123. No knockdown of phospho-AKT (Ser473) or phospho-PRAS40 (Y246) indicating specificity of siRNAs for human AKT.

**Supplemental Figure 4:**
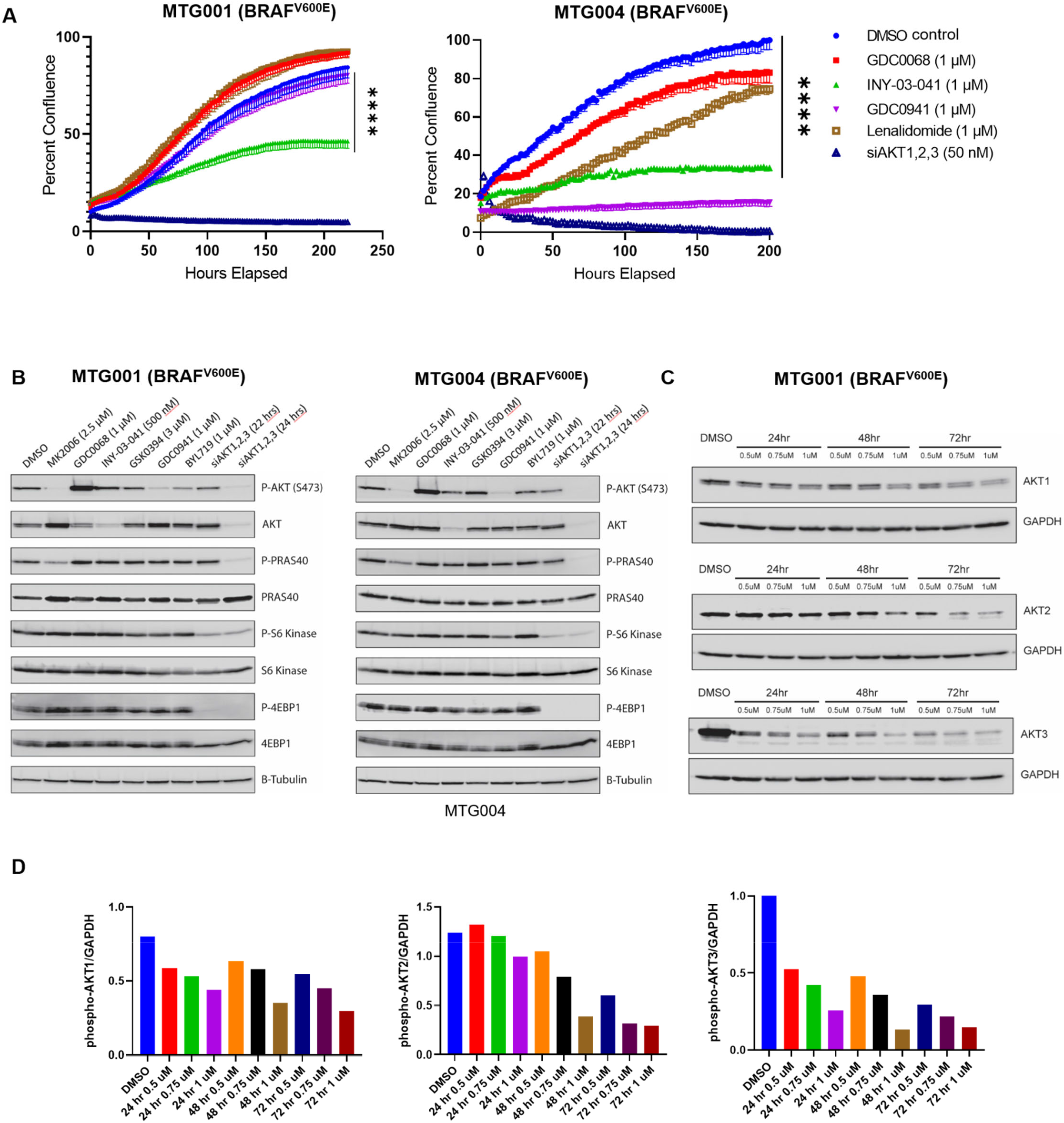
INY-03-041 leads to substantial effect on melanoma cell proliferation but not as potent as siAKT123. A) Cell confluency assay of MTG001 and MTG004 cell lines treated with DMSO control, GDC0068 (1μM), INY-03-041 (1μM), GDC0941 (1μM), Lenalidomide (1μM), and siAKT123 (50nM). Treatment with INY-03-041 lead to decreased cell proliferation (p<0.0001 in each cell line) but did not lead to cell death as with siAKT123 in MTG001 and MTG004. Lenalidomide, recruiter of the E3 ubiquitin ligase Cereblon and conjugated to GDC0068 to form INY-03-041, was used as a negative control for off-target effects of INY-03-041. Error bars indicate standard error of the mean of triplicate wells. B) Immunoblotting of cell lysates treated with DMSO control, MK2206 (2.5μM), GDC0068 (1μM), INY-03-041 (500nM), GSK0394 (3μM), GDC0941 (1μM), BYL719 (1μM), siAKT123 (50nM) at 22 hours, and siAKT123 (50nM) at 24 hours reveals complete knockdown of pAKT (Ser473) and total AKT by siAKT123 at 24 hours, as well as knockdown of p-PRAS40, p-S6 Kinase, and p-4EBP1. INY-03-041 displays 77% reduction of total AKT compared to 89% reduction by siAKT123. C) Time-course immunoblotting of cell lysates treated with varying concentrations of INY-03-041 demonstrated incomplete total protein knockdown of individual AKT paralogs. D) Densitometry quantification of time-course immunoblotting found in (B).

**Supplemental Figure 5:**
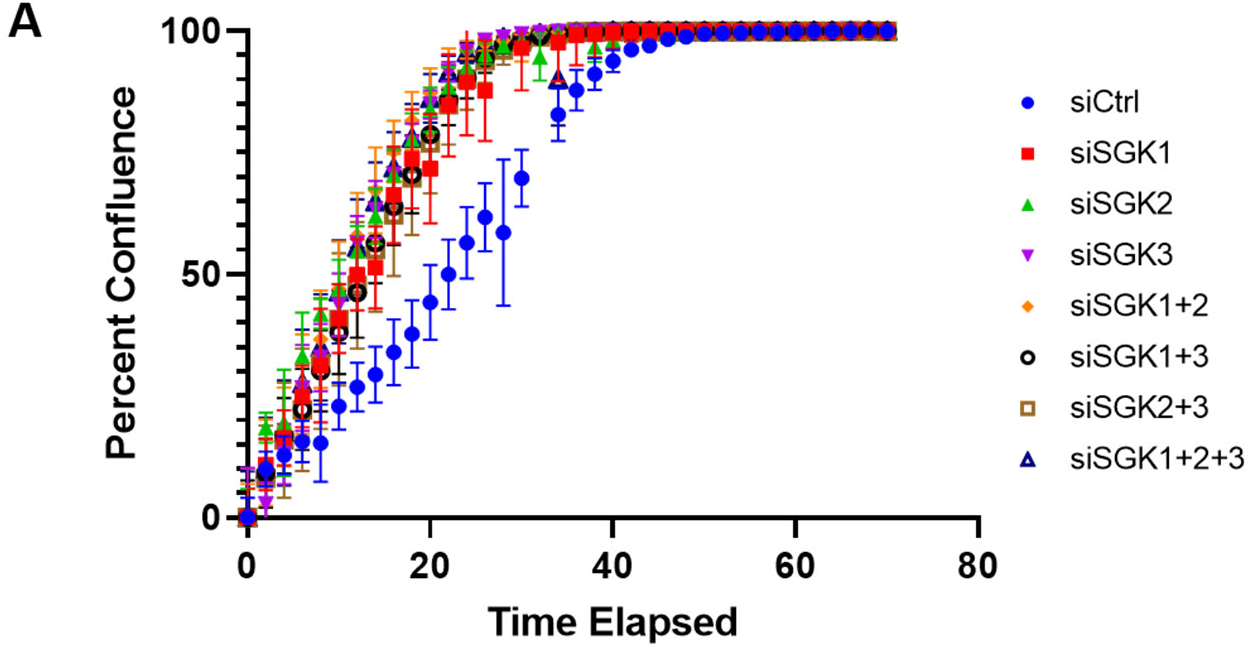
SiRNA-mediated knockdown of SGK does not inhibit melanoma cell viability. A) Cell confluency analysis of A375 cells treated with siCtrl, siSGK1, siSGK2, siSGK3, siSGK1+2, siSGK1+3, siSGK2+3, and siSGK1+2+3 demonstrated no effect with any combination of siRNAs.

**Supplemental Figure 6:**
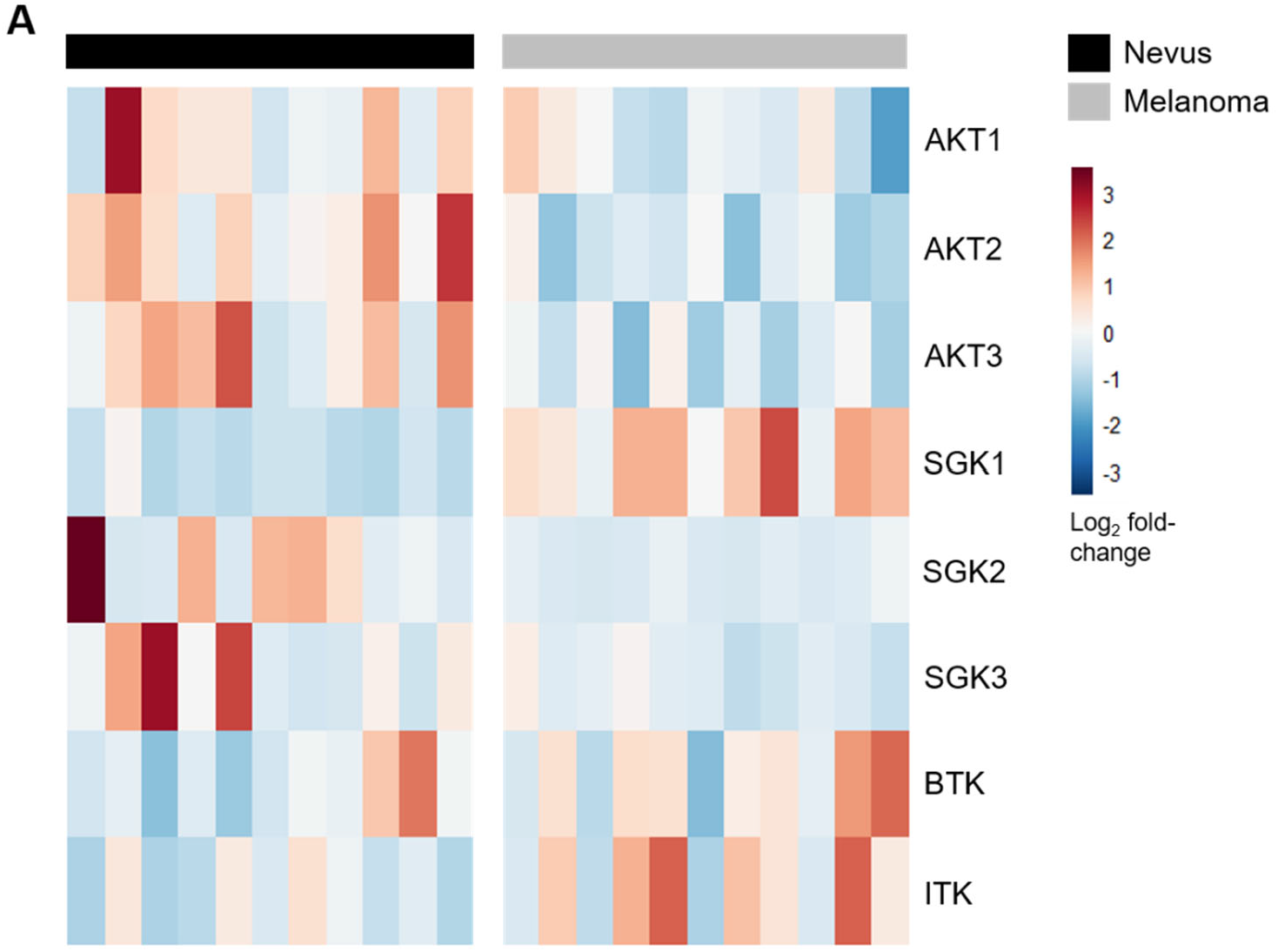
SGK1 is significantly upregulated in melanoma vs. nevus. A) Differential expression analysis from RNA sequencing of matched nevi vs primary melanoma samples revealed that SGK1 is upregulated in melanoma (p=0.0001).

## METHODS

### Viral Constructs and Propagation

The avian retroviral vectors used in this study are replication-competent Avian Leukosis Virus splice acceptor and Bryan polymerase-containing vectors of envelope subgroup A [designated RCASBP(A) and abbreviated RCAS]. A Gateway compatible RCAS destination vector has been described (43). RCAS-Cre (44) and RCAS-myrAKT1 also have been described (45,46). RCAS-myrAKT1 was used as a template to generate the *Akt1* phosphorylated mutant constructs using Q5® Site-Directed Mutagenesis (New England Biolabs). Myr-Akt was mutated with the following primers: S473A FWD “CACTTCCCCCAGTTCGCCTACTCAGCCAGTGGC”, S473A REV “GCCACTGGCTGAGTAGGCGAACTGGGGGAAGTG”, T308A FWD “TGCCACTATGAAGGCATTCTGCGGAACGCCG”, T308A REV “CGGCGTTCCGCAGAATGCCTTCATAGTGGCA.” S473D FWD “CACTTCCCCCAGTTCGACTACTCAGCCAGTGGC”, S473D REV “GCCACTGGCTGAGTAGTCGAACTGGGGGAAGTG”, T308D FWD “TGCCACTATGAAGGATTTCTGCGGAACGCCG,” and T308D REV “CGGCGTTCCGCAGAAATCCTTCATAGTGGCA.” pDONR223-SGK (wildtype) was obtained from Addgene and recombined into the destination vector pDEST FG12_CMV_GW, SV40_Luciferase-IRES-eGFP. pDONR223-SGK used as a template to generate the myr SGK1 construct as well as SGK1 kinase dead (K127M) constructs using Site-Directed Mutagenesis (New England Biolabs). SGK1 was mutated with the following primers: K127M FWD: CTATGCAGTCATGGTTTTACAGAAGAAAG, K127M REV: AACACTTCTTCTGCCTTG.

### Cell Culture

Melanoma cell lines A375, HT144, SK-MEL28, WM793, YUMM3.2 (Yale University Mouse Melanoma 3.2) cells were cultured in DMEM/F12 media (Thermo Fisher) supplemented with 10% FBS (Atlanta Biologicals, Flowery Branch, GA) and maintained at 37°C. DF-1 cells were grown in DMEM-high glucose media (Thermo Fisher) supplemented with 10% FBS (Atlas Biologicals) and 0.5 μg/mL Gentamicin (Thermo Fisher), and maintained at 39°C.

### Viral Infections (*In vitro*)

A375, HT144, and WM793 cells were transfected with pcDNA3.1-*TVA* (47) containing the hygromycin B-resistance gene to generate A375-TVA, HT144-TVA, and WM793-TVA cells. TVA-positive clones were selected in 300 μg/mL Hygromycin B (Thermo Fisher). Supernatant from DF-1 cells producing RCAS myr-HA-Akt1-T308A; RCAS myr-HA-Akt1-T308D; RCAS myr-HA-Akt1-S473A; myr-HA-Akt1-S473D; myr-HA-Akt1-T308A, S473A; myr-HA-Akt1-T308A, S473D; myr-HA-Akt1-T308D, S473A; myr-HA-Akt1-T308D, S473D; myr-HA-Akt1-delta(PH); and myr-HA-Akt1-delta(PH)-K179M was used to infect A375, HT144, and WM793 TVA+ cells. Expression of Akt1 mutants in A375-TVA+, HT144-TVA+, and WM793-TVA+ cells were confirmed by immunoblot for HA. To generate the YUMM 3.2 SGK1 isogenic cell lines, myr-SGK1 was gateway cloned from pDONR221 TOPO to a FG12 CMV--Luciferase-IRES-eGFP destination vector to generate FG12 CMV myr-SGK1;Luciferase-IRES-eGFP. The lentiviral vector FG12 CMV myr-SGK1;Luciferase-IRES-eGFP, along with packaging plasmids psPAX2 (#12260 Addgene, Cambridge, MA) and pCMV-VSV-G (#8454 Addgene) were transfected into 293FT cells using the calcium phosphate method. Supernatant from these cells containing virus was then used to infect isogenic YUMM 3.2 parental and Pten^-/-^ cell lines YUMM3.2 SGK1 cells were sorted for eGFP using a Propel Labs Avalon at The Flow Cytometry Shared Resource Laboratory at the University of Utah.

### Pharmacological Inhibitors

The AKT inhibitors (MK2206, GDC690693, GSK-2141795), PI3K inhibitors (GDC0941, BYL719) and SGK inhibitor (GSK690394) were purchased from Selleck Chemical (Houston, TX). GDC-0068 was generously provided by Genentech. The AKT PROTAC, INY-03-041, was kindly provided by Dr. Nathanael S. Gray (Stanford University) and Dr. Alex Toker (Harvard Medical School). Each drug was formulated in DMSO and added to F12/DMEM (Thermo Fisher) media for a final concentration of 0.1% (v/v) DMSO. Inhibitors were used at select concentrations as indicated in figures and text.

### Immunoblotting

Protein extracts were harvested by lysing cells in radioimmunoprecipitation assay (RIPA) lysis buffer containing Halt Protease and Phosphotase Inhibitor Cocktail (Thermo Scientific). A BCA assay was performed to determine protein concentration prior to boiling. Samples were standardized to equal concentrations and diluted into NuPAGE LDS (lithium dodecyl sulfate, pH 8.4) sample loading buffer (Invitrogen) prior to boiling for 10 minutes at 95°C. Proteins were separated using standard SDS-PAGE gel electrophoresis with 8-16% gradient Tris-Glycine polyacrylamide gels, transferred to nitrocellulose membranes for immunoblot analysis using an iBlot2 (Invitrogen), and stained with primary antibodies as indicated in each figure. LI-COR brand Intercept (TBS) blocking buffer and IRDye RD and 800CW secondary antibodies were utilized for all experiments. Membranes were scanned and analyzed using the Odyssey Imaging System and software (LI-COR). Antibodies used: HA (MMS-101P; Biolegend), AKT (4691; Cell Signaling Technology), Phospho-AKT (T308) (13038; Cell Signaling Technology), Phospho-AKT (S473) (3787; Cell Signaling Technology), PRAS40 (2691; Cell Signaling Technology), Phospho-PRAS40 (Y246) (2997; Cell Signaling Technology), PTEN (9188; Cell Signaling Technology), Phospho-p44/42 MAPK (4370; Cell Signaling Technology), p44/42 MAPK (9102; Cell Signaling Technology), Phospho-S6 ribosomal protein (Ser235/235) (4858; Cell Signaling Technology), S6 ribosomal protein (2217; Cell Signaling Technology), Akt1 (2938; Cell Signaling Technology), Akt2 (3063, Cell Signaling Technology), Akt3 (3788; Cell Signaling Technology), Phospho-4E-BP1 (T37/46) (2855; Cell Signaling Technology), 4E-BP1 (9644; Cell Signaling Technology); SGK1 (D72C11; Cell Signaling Technology); Phospho-NDRG1 (Thr348) (D98G11; Cell Signaling Technology); NDRG1 (D8G9; Cell Signaling Technology).

### Incucyte Live Cell Proliferation Assays

Cell proliferation was assessed by seeding ~3,000 cells (A375, WM793), ~5,000 cells (HT144, SK-MEL28), or ~7,000 cells (MTG1, MTG4) per well in 96-well plates. Experiments were performed on two-to-three separate occasions in triplicate wells per experimental condition. Pharmacological agents were added 24 hours after cel plating. Cells were cultured in the absence or presence of pharmacological agents for 3-5 days or until the controls reached ~100% confluency. Confluence was assessed over time using an IncuCyte® Zoom Live Cell Imaging instrument with data analyzed using IncuCyte® Analysis Software (Sartorius) in two-hour intervals.

### Luminex Assay

Luminescence measurements were carried on a Millipore MAGPIX® according to the manufacturer’s instructions, using the following kit: Bio-Plex Pro Cell Signaling Akt panel, 8-plex (lq00006jk0k0rr).

### Lipofectamine-mediated siRNA transfection

siRNAs against human AKT1 (J-003000-10-0005 and J-003000-11-0005), AKT2 (J-003001-09-0005 and J003001-10-0005), AKT3 (J-003002-13-0005 and J-003002-14-0005), SGK1 (J-003027-13-0005), SGK2 (J-004673-09-0005), and SGK3 (J-004162-06-0005) genes and a negative control siRNA with scramble sequence were purchased from Dharmacon (and transfected with Lipofectamine RNAiMAX reagent at a total concentration of 50nM per siRNA-treated condition. In order to test for rescue of the effects of siAKT1-3, constructs encoding mouse AKT protein kinases, which are resistant to the human-specific siRNAs described above, were generated and expressed in human melanoma cells with expression confirmed by immunoblotting. Rescue assays were performed in media supplemented with 10%(v/v) FBS as previously described using Cell Titer 96 MTS (Promega). Briefly, cells were seeded at ~50% confluency the day prior to transfection. Lipofectamine-mediated siAKT1-3 transfection (as described above) was performed at time point 0, and an MTS assay was performed 48 hours post-transfection. Each assay was performed in triplicate with at least 3 independent technical replicates. Data was plotted and analyzed using GraphPad Prism Software.

### Phospho-PRAS40 enzyme-linked immunosorbent assays (ELISA)

Levels of phosphorylated proline-rich AKT substrate of 40kDa (phospho-PRAS40 (pT246)) were assessed using PRAS40 ELISA kits (Invitrogen, Camarillo, USA, KHO0421). Cells were lysed in extraction buffer (Invitrogen, Camarillo, USA, FNN0011), and samples were diluted 1 in 50 in diluent buffer. Absorbance was measured at 450nm upon completion of assay.

### *In vivo* tumorigenesis studies

Four to eight-week-old immune-competent mice tolerized to EGFP and luciferase were injected subcutaneously into the right flank with 2.5×10^5^ melanoma cells and observed for tumorigenic growth. Tumors were visualized and measured weekly using bioluminescence imaging, and tumor burden was quantified using luminoscore values and digital caliper measurements (Day 2014). The following formula was used to calculate tumor volume: (Length x Width^2^)/2. All mouse tissues were fixed in 10% neutral buffered formalin overnight, then dehydrated in 70% ethyl alcohol.

### Bioluminescence Imaging

Mice were injected intraperitoneally with 16.7 mg/mL D-Luciferin in 200μL of phosphate buffered saline 10 minutes prior to image acquisition. The IVIS Spectrum was used to acquire images at one-week intervals beginning one-week after post-subcutaneous implantation of mouse YUMM3.2 melanoma cells until the experimental endpoint (6 weeks post-injection) or when tumor burden reached 2000mm^3^. Living Image software (version 4.5.2) was used to compile images.

### Statistical Methods

Mouse censored survival data was analyzed using a log-rank (Mantel-Cox) test of the Kaplan-Meier estimate of survival; a Fisher’s exact test was used to determine differences in the incidence of metastasis between cohorts. Densitometry measurements were performed using ImageJ and protein levels were normalized to GAPDH; the data are represented as mean ± S.E.M.

### Study Approval

All animal experimentation was performed in AAALAC approved facilities at the University of Utah. All animal protocols were reviewed and approved prior to experimentation by the Institutional Animal Care and Use Committee (IACUC) at the University of Utah.

## REFERENCES

1. Ascierto PA, Kirkwood JM, Grob JJ, Simeone E, Grimaldi AM, Maio M, et al. The role of BRAF V600 mutation in melanoma. J Transl Med 2012;10:85

2. Eroglu Z, Ribas A. Combination therapy with BRAF and MEK inhibitors for melanoma: latest evidence and place in therapy. Ther Adv Med Oncol 2016;8:48–56

3. Damsky W, Micevic G, Meeth K, Muthusamy V, Curley DP, Santhanakrishnan M, et al. mTORC1 activation blocks BrafV600E-induced growth arrest but is insufficient for melanoma formation. Cancer Cell 2015;27:41–56

4. Dankort D, Curley DP, Cartlidge RA, Nelson B, Karnezis AN, Damsky WE, Jr., et al. Braf(V600E) cooperates with Pten loss to induce metastatic melanoma. Nat Genet 2009;41:544–52

5. Davies MA. The role of the PI3K-AKT pathway in melanoma. Cancer J 2012;18:142–7

6. Deuker MM, McMahon M. Rational targeting of BRAF and PI3-Kinase signaling for melanoma therapy. Mol Cell Oncol 2016;3:e1033095

7. Manca A, Lissia A, Capone M, Ascierto PA, Botti G, Caraco C, et al. Activating PIK3CA mutations coexist with BRAF or NRAS mutations in a limited fraction of melanomas. J Transl Med 2015;13:37

8. Sanchez-Vega F, Mina M, Armenia J, Chatila WK, Luna A, La KC, et al. Oncogenic Signaling Pathways in The Cancer Genome Atlas. Cell 2018;173:321–37 e10

9. Kircher DA, Trombetti KA, Silvis MR, Parkman GL, Fischer GM, Angel SN, et al. AKT1(E17K) Activates Focal Adhesion Kinase and Promotes Melanoma Brain Metastasis. Mol Cancer Res 2019;17:1787–800

10. Mahajan K, Mahajan NP. PI3K-independent AKT activation in cancers: a treasure trove for novel therapeutics. J Cell Physiol 2012;227:3178–84

11. Martelli AM, Tabellini G, Bressanin D, Ognibene A, Goto K, Cocco L, et al. The emerging multiple roles of nuclear Akt. Biochim Biophys Acta 2012;1823:2168–78

12. Saji M, Ringel MD. The PI3K-Akt-mTOR pathway in initiation and progression of thyroid tumors. Mol Cell Endocrinol 2010;321:20–8

13. Dai DL, Martinka M, Li G. Prognostic significance of activated Akt expression in melanoma: a clinicopathologic study of 292 cases. J Clin Oncol 2005;23:1473–82

14. Durban VM, Deuker MM, Bosenberg MW, Phillips W, McMahon M. Differential AKT dependency displayed by mouse models of BRAF(V600E)-initiated melanoma. J Clin Invest 2013;123:5104–18

15. Dinavahi SS, Noory MA, Gowda R, Drabick JJ, Berg A, Neves RI, et al. Moving Synergistically Acting Drug Combinations to the Clinic by Comparing Sequential versus Simultaneous Drug Administrations. Mol Pharmacol 2018;93:190–6

16. Rozengurt E, Soares HP, Sinnet-Smith J. Suppression of Feedback Loops Mediated by PI3K/mTOR Induces Multiple Overactivation of Compensatory Pathways: An Unintended Consequence Leading to Drug Resistance. Mol Cancer Ther 2014;13:2477–88

17. Silva JM, Bulman C, McMahon M. BRAFV600E cooperates with PI3K signaling, independent of AKT, to regulate melanoma cell proliferation. Mol Cancer Res 2014;12:447–63

18. Di Cristofano A. SGK1: The Dark Side of PI3K Signaling. Curr Top Dev Biol 2017;123:49–71

19. Study of PX-866 and Vemurafenib in Patients With Advanced Melanoma. <https://clinicaltrials.gov/ct2/show/NCT01616199>.

20. BKM120 Combined With Vemurafenib (PLX4032) in BRAFV600E/K Mutant Advanced Melanoma.

21. Mundi PS, Sachdev J, McCourt C, Kalinsky K. AKT in cancer: new molecular insights and advances in drug development. Br J Clin Pharmacol 2016;82:943–56

22. Lazaro G, Kostaras E, Vivanco I. Inhibitors in AKTion: ATP-competitive vs allosteric. Biochem Soc Trans 2020;48:933–43

23. Hirai H, Sootome H, Nakatsuru Y, Miyama K, Taguchi S, Tsujioka K, et al. MK-2206, an allosteric Akt inhibitor, enhances antitumor efficacy by standard chemotherapeutic agents or molecular targeted drugs in vitro and in vivo. Mol Cancer Ther 2010;9:1956–67

24. Sun L, Huang Y, Liu Y, Zhao Y, He X, Zhang L, et al. Ipatasertib, a novel Akt inhibitor, induces transcription factor FoxO3a and NF-kappaB directly regulates PUMA-dependent apoptosis. Cell Death Dis 2018;9:911

25. You I, Erickson EC, Donovan KA, Eleuteri NA, Fischer ES, Gray NS, et al. Discovery of an AKT Degrader with Prolonged Inhibition of Downstream Signaling. Cell Chem Biol 2020;27:66–73 e7

26. Vivanco I, Chen ZC, Tanos B, Oldrini B, Hsieh WY, Yannuzzi N, et al. A kinase-independent function of AKT promotes cancer cell survival. Elife 2014;3

27. Hart JR, Vogt PK. Phosphorylation of AKT: a mutational analysis. Oncotarget 2011;2:467–76

28. Manning BD, Toker A. AKT/PKB Signaling: Navigating the Network. Cell 2017;169:381–405

29. Hong F, Larrea MD, Doughty C, Kwiatkowski DJ, Squillace R, Slingerland JM. mTOR-raptor binds and activates SGK1 to regulate p27 phosphorylation. Mol Cell 2008;30:701–11

30. Sherk AB, Frigo DE, Schnackenberg CG, Bray JD, Laping NJ, Trizna W, et al. Development of a smallmolecule serum-and glucocorticoid-regulated kinase-1 antagonist and its evaluation as a prostate cancer therapeutic. Cancer Res 2008;68:7475–83

31. Parkman GL, Foth M, Kircher DA, Holmen SL, McMahon M. The role of PI3’-lipid signalling in melanoma initiation, progression and maintenance. Exp Dermatol 2022;31:43–56

32. Tolcher AW, Patnaik A, Papadopoulos KP, Rasco DW, Becerra CR, Allred AJ, et al. Phase I study of the MEK inhibitor trametinib in combination with the AKT inhibitor afuresertib in patients with solid tumors and multiple myeloma. Cancer Chemother Pharmacol 2015;75:183–9

33. Shoushtari AN, Ong LT, Schoder H, Singh-Kandah S, Abbate KT, Postow MA, et al. A phase 2 trial of everolimus and pasireotide long-acting release in patients with metastatic uveal melanoma. Melanoma Res 2016;26:272–7

34. Sweeney C, Bracarda S, Sternberg CN, Chi KN, Olmos D, Sandhu S, et al. Ipatasertib plus abiraterone and prednisolone in metastatic castration-resistant prostate cancer (IPATential150): a multicentre, randomised, double-blind, phase 3 trial. Lancet 2021;398:131–42

35. Vanhaesebroeck B, Perry MWD, Brown JR, Andre F, Okkenhaug K. PI3K inhibitors are finally coming of age. Nat Rev Drug Discov 2021;20:741–69

36. Alessi DR, James SR, Downes CP, Holmes AB, Gaffney PR, Reese CB, et al. Characterization of a 3-phosphoinositide-dependent protein kinase which phosphorylates and activates protein kinase Balpha. Curr Biol 1997;7:261–9

37. Alessi DR, Andjelkovic M, Caudwell B, Cron P, Morrice N, Cohen P, et al. Mechanism of activation of protein kinase B by insulin and IGF-1. EMBO J 1996;15:6541–51

38. Scortegagna M, Lau E, Zhang T, Feng Y, Sereduk C, Yin H, et al. PDK1 and SGK3 Contribute to the Growth of BRAF-Mutant Melanomas and Are Potential Therapeutic Targets. Cancer Res 2015;75:1399–412

39. Karbowniczek M, Spittle CS, Morrison T, Wu H, Henske EP. mTOR is activated in the majority of malignant melanomas. J Invest Dermatol 2008;128:980–7

40. Mendoza MC, Er EE, Blenis J. The Ras-ERK and PI3K-mTOR pathways: cross-talk and compensation. Trends Biochem Sci 2011;36:320–8

41. Castel P, Scaltriti M. The emerging role of serum/glucocorticoid-regulated kinases in cancer. Cell Cycle 2017;16:5–6

42. Toker A. mTOR and Akt signaling in cancer: SGK cycles in. Mol Cell 2008;31:6–8

43. Loftus SK, Larson DM, Watkins-Chow D, Church DM, Pavan WJ. Generation of RCAS vectors useful for functional genomic analyses. DNA Res 2001;8:221–6

44. VanBrocklin MW, Robinson JP, Lastwika KJ, Khoury JD, Holmen SL. Targeted delivery of NRASQ61R and Cre-recombinase to post-natal melanocytes induces melanoma in Ink4a/Arflox/lox mice. Pigment Cell & Melanoma Research 2010;23:531–41

45. Aoki M, Batista O, Bellacosa A, Tsichlis P, Vogt PK. The akt kinase: molecular determinants of oncogenicity. Proceedings of the National Academy of Sciences of the United States of America 1998;95:14950–5

46. Cho JH, Robinson JP, Arave RA, Burnett WJ, Kircher DA, Chen G, et al. AKT1 Activation Promotes Development of Melanoma Metastases. Cell reports 2015;13:898–905

47. Bromberg-White JL, Webb CP, Patacsil VS, Miranti CK, Williams BO, Holmen SL. Delivery of short hairpin RNA sequences by using a replication-competent avian retroviral vector. J Virol 2004;78:4914–6

